# The trade-off between parsimony and model complexity for understanding biomedical mechanisms from mathematical models

**DOI:** 10.64898/2026.08.31.748397

**Authors:** Patricia Lamirande, Mia Brunetti, Terry Easlick, Fatemeh Beigmohammadi, Morgan Craig

## Abstract

Mechanistic mathematical models have been used extensively to provide a deeper understanding of biological mechanisms, including unveiling the regulation of tumour growth and its response to various treatments. However, given the breadth of biological regulatory mechanisms, these models are frequently large and thus prone to potential issues with parameter identifiability. Statistical metrics like the Akaike and Bayesian information criteria can help identify a parsimonious model by balancing goodness of fit against model complexity. Yet simple models may fail to provide sufficient biological insight if they do not adequately capture known physiological processes or mechanisms. A modeller must therefore balance hypothesis generation and biological learning with model tractability. Here, we illustrate this balance using models of ovarian cancer growth and treatment response to cisplatin and immune checkpoint blockade in homologous recombination (HR)-deficient and HR-proficient immunocompetent mouse models. We develop a hierarchy of mathematical models of increasing complexity to describe tumour growth, treatment response, and immune dynamics. Our results highlight the limits of relying purely on statistical metrics for model selection, particularly when the goal is to obtain biological insight and underscore the importance of balancing model complexity to avoid overfitting and parameter unidentifiability.

## Introduction

Ovarian cancer is largely diagnosed at an advanced stage^1^. This late diagnosis coupled with the fact that ovarian cancers will unavoidably become resistant to standard-of-care platinum-based chemotherapy^2^ place them among the most lethal of cancers^1^ and a leading cause of female cancer death^3^. Immunotherapies, including immune checkpoint blockade (ICB), are now a standard component of anti-cancer treatments^4^. For many patients, ICB (alone or in combination) provides effective and durable responses^5^. Unfortunately, ICB has failed to meet efficacy endpoints in ovarian cancer clinical trials^6^. Several factors contribute to these disappointing results, including a general lack of predictive biomarkers^7^, an incomplete understanding of the mechanisms of response^4^ for ICB, and the impact of heterogeneity in the ovarian cancer tumour microenvironment on treatment response^1,8^. Further, preclinical and animal models of many ovarian cancers do not exist^9,10^ or are insufficient^11^, impeding drug development. In response, Paffenholz et al.^11^ developed mouse models of highgrade serous ovarian cancer (HGSOC) to better understand therapeutic responses to platinum-based chemotherapy (i.e., cisplatin), ICB, and their combinations in homologous recombination (HR)-deficient and proficient immunocompetent mice, called MP and MPB1 respectively. Identifying and interpreting mechanisms of response in these different scenarios can be difficult due to the multiple overlapping and nonlinear tumour, immune, and drug responses. Further, translating preclinical findings to patients is complicated by a host of factors whether they are biological (e.g., degree of heterogeneity in the tumour microenvironment of ovarian cancers^1,8^, differences in subtypes, etc.) or drug-related (e.g., number of possible treatment schedules, including potential orderings). These make evaluating all possible combinations for safety and efficacy especially difficult. To this end, mathematical models incorporating key disease phenotypes and treatment effects can be used to rationalize the scheduling of treatment combinations.

Developing a mathematical model requires striking a balance between biological realism and model tractability. This balance is particularly critical when approaching a multifactorial problem like developing new treatment approaches in a disease as complex as ovarian cancer. Modellers use priors (statistical, biological, or otherwise) that influence model design; some may prefer the simplest possible model, whereas others may prefer more complex models for various reasons. Several techniques exist to evaluate how to weigh model simplicity with biological interpretability and significance. These include statistical parsimony measures like the Akaike and Bayesian information criteria (AIC and BIC, respectively), sensitivity analyses, and structural and practical parameter identifiability techniques. In contrast to statistically based models (e.g., machine learning/artificial intelligence, general linear models, nonlinear mixed effects models, etc.), mechanistic models provide causal links between their parameters and observed dynamics^12^. In certain disciplines, like quantitative systems pharmacology, such mechanistic models can encompass tens-to-hundreds of equations and hundreds of parameters. Thus, to deal with parameter identifiability, many parameters are tightly constrained, often fixed to constant values, while key parameters may be allowed to range more widely through estimation from data. This raises two key questions: 1) what information is lost, if any, if we uniquely favour the selection of the most parsimonious model and 2) are we in danger of “over-modelling” and losing confidence in our predictions by using too large a model? In the former case, over-reliance on maximizing parsimony can lead to erroneous conclusions (the “long-branch attraction” problem^13,14^ in genetics is one such example). In the latter case, we can have little confidence in the predictions of an over-fitted or over-parameterized model.

The tension between model simplicity and biological realism has long been recognized in mathematical oncology. For example, Kohandel, Sivaloganathan, and Oza^15^ compared multiple formulations describing tumour-growth and treatment-responses, emphasizing that the purpose of mathematical modelling is not merely to reproduce observed data, but to understand how biological hypotheses are translated into quantitative predictions. Motivated by this philosophy, we developed several models of expanding complexity describing ovarian cancer growth and treatment with cisplatin and ICB, based on the results of Paffenholz et al.^11^, to explore what can be learned by increasingly complex mechanistic models of cancer progression and treatment. Our main interest was to gain insight into the consequences of progressively adding additional mechanisms on model fidelity to data and the degree of biological insight gained. Comparing model performance across genetic and treatment contexts showed that multiple levels of biological detail reproduced tumour growth while revealing distinct biological differences between MP and MPB1 models (i.e., HR-proficient and HR-deficient tumours). Our findings suggest that the improved therapeutic response of HR-deficient tumours arise from both enhanced intrinsic sensitivity to cisplatin and a more favourable anti-tumoural immune response. More broadly, our modelling framework demonstrates that increasing model complexity can provide mechanistic insight beyond that obtained from simpler tumour growth models.

## Methods

### Mathematical models of tumour growth

To assess the impact of incorporating additional complexity within mechanistic modelling approaches, we developed various mathematical models to describe the growth of ovarian cancer tumours in MP and MPB1 mice without treatment and when treated with cisplatin, ICB, and a combination of both.

In the simplest model, we used standard tumour growth models^16^ and ignored tumour-immune interactions and the pharmacokinetics/pharmacodynamics (PK/PD) of both cisplatin and ICB. Let *g*(*T*(*t*)) describe tumour growth, where *T*(*t*) is the tumour volume (in mm^3^) at time *t*. The exponential, logistic, and Gompertz growth models are given by

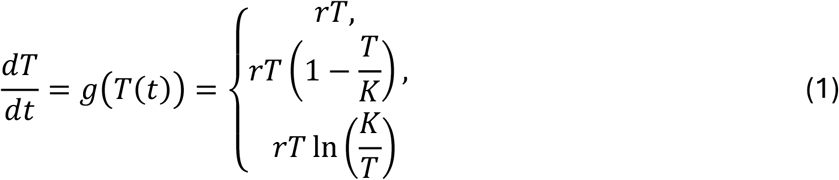

respectively, where *r* represents the tumour’s intrinsic growth rate (in days^−1^), and *K* denotes the tumour’s carrying capacity (in mm^3^).

To add a second level of complexity, we incorporated in Eq. (1) a PK/PD model to describe cisplatin exposure and its effects on tumour growth (**Figure 1B**). This was done by integrating cisplatin exposure based on the population PK model developed by Urien and Lokiec^17^. Let *C*_1_ and *C*_2_ (in μg*⋅L*^−1^) be the concentration of cisplatin in the central (i.e., plasma) and peripheral compartments, respectively, and *k*_*ij*_ (in day^−1^) denote the transit rate from compartment *i* to compartment *j* (for *i* = 1,2; *j* = 0,1,2). Then, the pharmacokinetics of cisplatin can be described by

**Figure 1.**
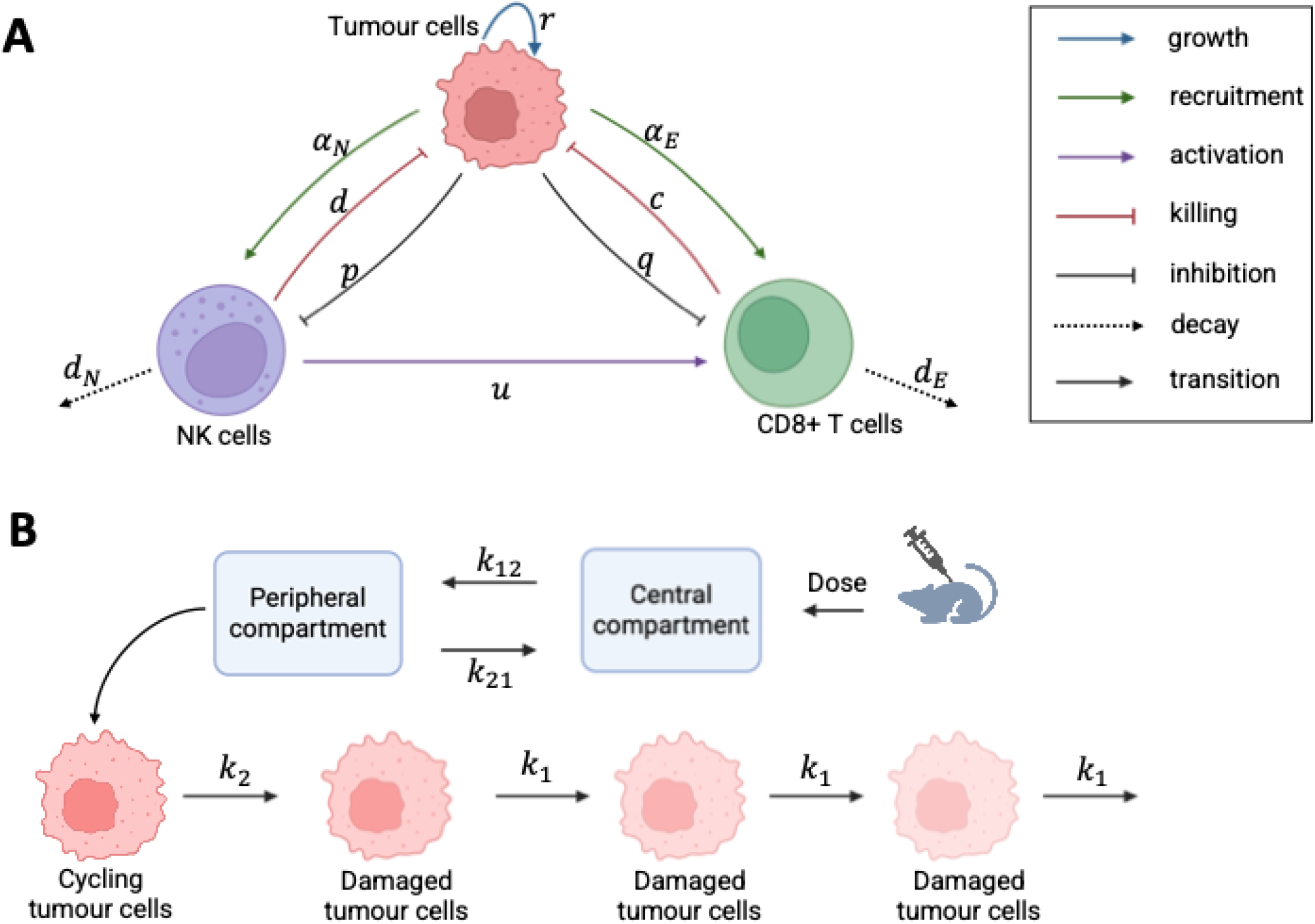
Model schematics. **A)** Graphical description of the mathematical models in Eqs. (1), (4), and (5) describing tumour growth without and with the immune response from NK and CD8+ T cells. **B)** Graphical description of the cisplatin PK/PD models described in Eqs. (2) and (3).

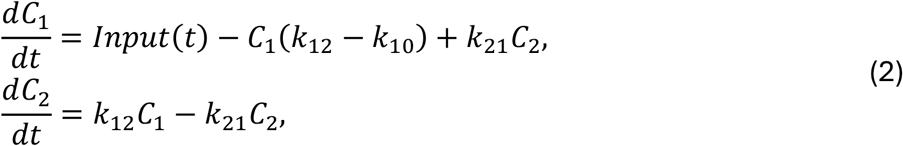

where *Input*(*t*) models the administration of the cisplatin dose^17^. In Paffenholz et al.^11^, cisplatin was administered by bolus injection, which we modelled as:

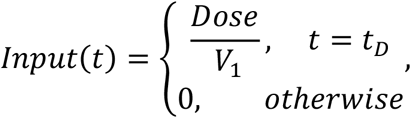

where *Dose* is the amount of cisplatin (μg), *t*_*D*_ (days) is the time the dose is administered, and *V*_*1*_ (L) is the volume of the central compartment. Cisplatin PDs were modelled according to Simeoni et al.^18^ who developed a transit compartment model to represent the delayed onset of the drug’s effects on tumour growth. Let *T*_1_ represent the volume of proliferating tumour cells, and *T*_*i*_(*t*) the volume of tumour cells in various states *i* (*i* = 2,3,4) of damage. The pharmacodynamic effects of cisplatin are modelled by the following ordinary differential equations model:

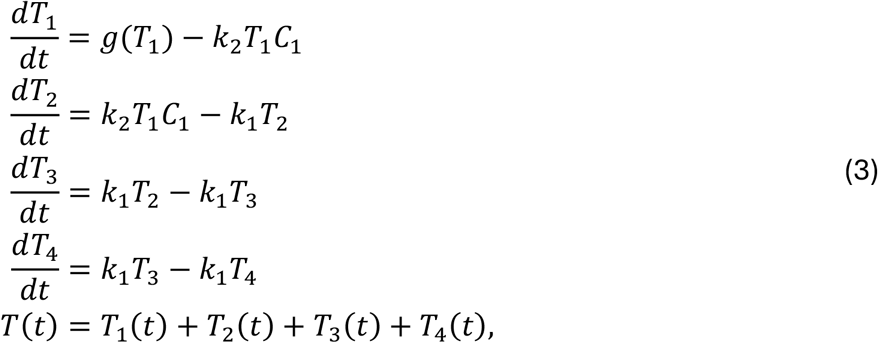

where *k*_1_ is the first-order transfer rate describing the kinetics of cell death (day^−1^) and *k*_2_ is the linear killing rate of cisplatin (L*⋅*mg^−1^day^−1^).

Given the importance of the tumour microenvironment and tumour-immune interactions on overall growth^8,19-23^, we added further complexity by incorporating tumour killing by interferon gamma (IFN-γ)-producing cells, namely natural killer (NK) and CD8+ T cells (**Figure 1A**). We drew on the model developed and validated by de Pillis et al.^24^ to describe tumour–immune interactions and the roles of NK cells and CD8+ T cells in controlling tumour growth. In their work, a simple killing term proportional to the NK cell population size effectively modelled tumour killing by NK cells. However, the same assumption did not accurately capture the action of CD8+ T cells. Therefore, for tumour-specific CD8+ T cells, they introduced a new killing function in which CD8+ T cells killing depends on the effector to target ratio and saturates at a maximum killing rate, resulting in an improved fit to the data. With *N* the number of NK cells and *E* the number of CD8+ T cells, these changes to Eq. (1) correspond to:

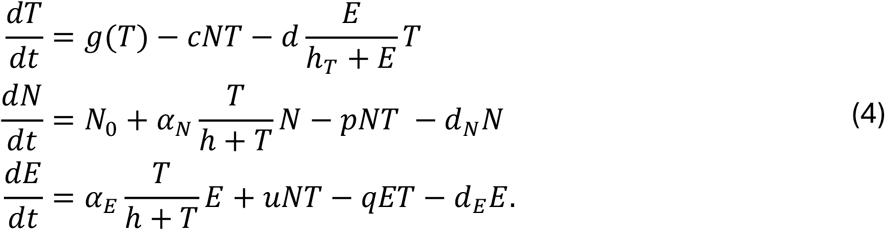

Here, *p* and *q* denote the rates of NK and CD8+ T cell death, respectively, through tumour cell interactions (cell^−1^day^−1^); all other parameters are defined in **Table 1**. In this model (Eq. (4), **Figure 1A**), both NK cells and CD8+T cells kill tumour cells, with both cell types eventually becoming inactivated due to their encounters with tumour cells.

**Table 1.** Fixed parameters values. Parameter names, descriptions, fixed values, and sources.

| Fixed parameters |  |  |  |
| --- | --- | --- | --- |
| Parameter | Description and units | Value | Reference |
| $\alpha_N$ | Maximal rate of NK cell activation by tumour cells (day<sup>-1</sup>) | $2.5 \times 10^{-2}$ | de Pillis et al. 2005 |
| $\alpha_E$ | Maximal rate of CD8+ T cell activation by tumour cells (day<sup>-1</sup>) | $3.75 \times 10^{-2}$ | de Pillis et al. 2005 |
| $c$ | Rate of tumour cell mass action killing by NK cells (cells<sup>-1</sup>·day<sup>-1</sup>) | $3.50 \times 10^{-6}$ | de Pillis et al. 2005 |
| $d_N$ | Natural death rate of NK cells (day<sup>-1</sup>) | $4.12 \times 10^{-2}$ | de Pillis et al. 2005 |
| $d_E$ | Natural death rate of CD8+ T cells (day<sup>-1</sup>) | $2 \times 10^{-2}$ | de Pillis et al. 2005 |
| $d$ | Maximal rate of tumour cell killing by CD8+ T cells (day<sup>-1</sup>) | 1.43 | de Pillis et al. 2005 |
| $h$ | Half-saturation constant in the immune cell recruitment functions, i.e. volume of the tumour cells eliciting 50% of the maximal stimulation of NK and T cell recruitment (mm<sup>3</sup>) | $4.49 \times 10^{-3}$ | Adapted from de Pillis et al. 2005 (assuming 1 mm<sup>3</sup> = 10<sup>6</sup> cells) |
| $h_T$ | Half-saturation constant in the tumour-killing function, i.e. number of CD8+ T cells producing 50% of maximum tumour cell killing (cells) | $1.18 \times 10^6$ | Mongeon and Craig 2025, assuming similar $E_0$ |
| $N_0$ | Constant rate of NK cell production (cells·day<sup>-1</sup>) | $1.3 \times 10^4$ | de Pillis et al. 2005 |
| $u$ | Stimulation rate of T cell to be produced because of tumour cells killed by NK cells (mm<sup>-3</sup> day<sup>-1</sup>) | 1.1$p$ | Based on 0.11 value in de Pillis et al. 2005 |

To simultaneously predict the effects of the tumour microenvironment and cisplatin on tumour cell growth, we combined the models in Eqs. (3) and (4) by modifying the tumour growth equation to account for the additional delayed killing of tumour cells by cisplatin. We further incorporated the increased death of immune cells induced by cisplatin, *k*_2_, and the drug’s recruitment of NK and CD8+ T cells in the microenvironment, with rates *r*_*C,N*_ and *r*_*C,E*_ respectively (L*⋅*mg^−1^day^−1^). These modifications give the system:

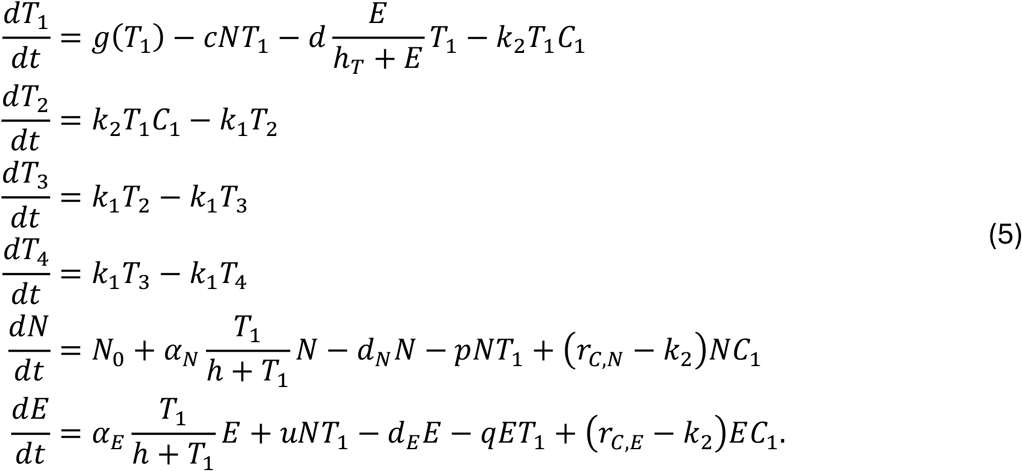

Lastly, we incorporated the effects of immunotherapy under ICB. Checkpoint proteins, such as PD-L1 on tumour cells and PD-1 on T cells, help keep immune responses in check. Cancer cells have evolved to express PD-L1, as the binding of PD-1 by PD-L1 prevents T cells from killing tumour cells. Immune checkpoint blockade (ICB) blocks this binding, allowing CD8+ T cells to kill tumour cells^7^. Previous models^25-29^ have described the mechanisms of immune checkpoint blockade explicitly by accounting for PD-1/PD-L1 binding dynamics and ICB pharmacokinetics. However, we instead opted to adjust parameter *q* in Eq. (4) to capture the ICB-induced change in tumour-mediated T-cell inactivation. This is because the necessary additional data to describe the more elaborate mechanisms behind ICB, such as immune cell counts under ICB treatment, was not measured in Paffenholz et al.^11^ Thus, introducing a fully mechanistic checkpoint model would substantially increase model complexity without sufficient information to constrain the additional parameters.

### Parameter estimation

To prevent overfitting and reduce the number of parameters fitted for each model (Eqs. (1), (3)-(5)), we fixed several parameter values based on previous modelling studies. Specifically, parameter estimates for equivalent terms in our model were obtained from the tumour-immune model of de Pillis et al.^24^ and the mechanistic model of glioblastoma response to ICB developed by Mongeon and Craig^25^. The full list of fixed parameters and their values is provided in **Table 1**.

We used a population pharmacokinetic study in mice performed by Fukushima et al.^30^ to fix the rate parameter values in the cisplatin PK model (Eq. (2)). Briefly, mice were administered 7.5 mg/kg of cisplatin via 30-second bolus injection and 2-hour infusion of 1.0, 2.5, 5.0, and 7.5 mg/kg, the latter with a total dosing volume of 6 mL. Following cisplatin administration, blood samples were collected 0.083, 0.25, 0.5, 0.75, 1, 1.5, and 2 hours after bolus injection, and 0.5, 1, 1.5, 2, 2.25, 2.5, 3, and 4 hours after infusion. A PK model equivalent to Eq. (3) was then fit to their data using a nonlinear mixed effects model, with bootstrapping of the data performed to calculate the 95% confidence interval about each mean parameter estimate and to assess the fidelity of the resulting parameter values. Mean values for the PK parameters in Eq. (3) are provided in **Table 2**. Note that transfer rate parameters *k*_*ij*_ were calculated according to the standard PK equation

**Table 2.**
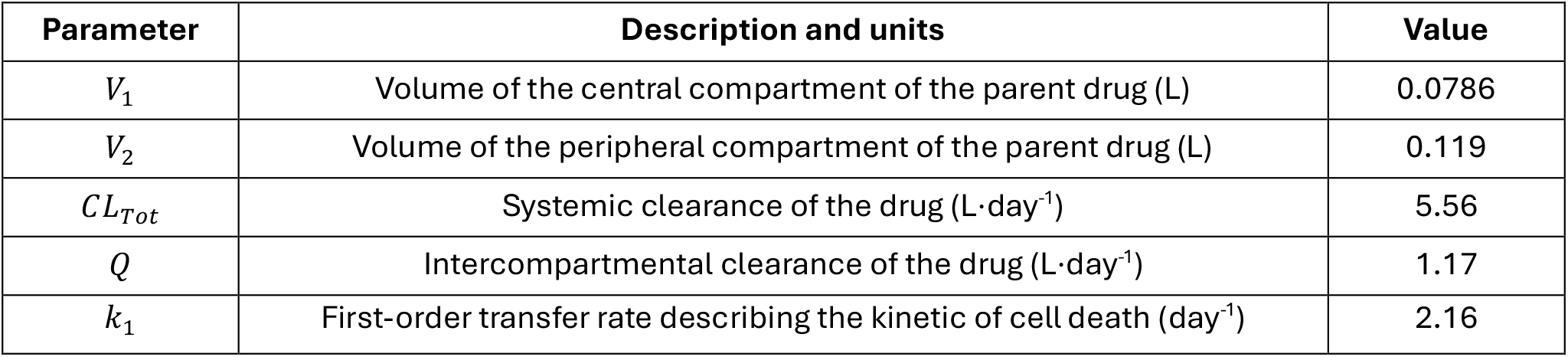
Parameter values of the cisplatin pharmacokinetic model. Selected cisplatin parameter value estimates from the original mouse data and bootstrap replicates of Fukushima et al.^30^ used in the cisplatin pharmacokinetic model (Eq. (2)).

| Parameter | Description and units | Value |
| --- | --- | --- |
| $V_1$ | Volume of the central compartment of the parent drug (L) | 0.0786 |
| $V_2$ | Volume of the peripheral compartment of the parent drug (L) | 0.119 |
| $CL_{Tot}$ | Systemic clearance of the drug (L·day<sup>-1</sup>) | 5.56 |
| $Q$ | Intercompartmental clearance of the drug (L·day<sup>-1</sup>) | 1.17 |
| $k_1$ | First-order transfer rate describing the kinetic of cell death (day<sup>-1</sup>) | 2.16 |

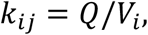

with the elimination rate *k*_10_ calculated as

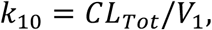

where *CL*_*tot*_ is the total rate of clearance.

The rest of the models’ parameters were estimated using data published by Paffenholz et al.^11^ In their study, they engineered two mouse models that support high-grade serous ovarian carcinomas; one HR-deficient with increased sensitivity to cisplatin, named MPB1, and the other HR-proficient, named MP. To gain mechanistic insight into genotype-dependent therapy responses to chemo- and immunotherapies, they measured tumour responses over time in both MP and MPB1 mice without treatment (vehicle), and treated with cisplatin chemotherapy, ICB, or the combination of the two. Paffenholz et al.^11^ also analyzed within-tumour NK and CD8+ T cells by flow cytometry. For this study, we used their longitudinal and flow cytometry data to calibrate and validate our models (see Figures 4C-D, 5A-B, and S5D in Paffenholz et al.^11^). See **Table 3** for the list of estimated parameters.

**Table 3.** Estimated parameters and their definitions. Estimated parameter values for each model are reported throughout the results and summarised in Table S5 (SI). Abbreviations: Veh, vehicle; Cis, cisplatin; ICB, immune checkpoint blockade.

| Parameter | Description and units | Estimated in |
| --- | --- | --- |
| $r$ | Intrinsic tumour growth rate (day<sup>-1</sup>) | Eq. (1): Veh, Cis, ICB, Cis + ICB; Eq. (4): Veh |
| $K$ | Tumour carrying capacity (mm<sup>3</sup>) | Eq. (1): Veh, Cis, ICB, Cis + ICB |
| $k_2$ | Linear killing rate of cisplatin (L·mg<sup>-1</sup>day<sup>-1</sup>) | Eq. (2)-(3): Cis |
| $p$ | Rate of NK cell inactivation by tumour cells (mm<sup>-3</sup> day<sup>-1</sup>) | Eq. (4): Veh |
| $q$ | Rate of CD8+T cells inactivation by tumour interaction (mm<sup>-3</sup> day<sup>-1</sup>) | Eq. (4): Veh; Eq. (4): ICB |
| $r_{C,N}$ | Recruitment of NK cells into TME due to cisplatin (L·mg<sup>-1</sup>day<sup>-1</sup>) | Eq. (5): Cis |
| $r_{C,E}$ | Recruitment of CD8+ T cells into TME due to cisplatin (L·mg<sup>-1</sup>day<sup>-1</sup>) | Eq. (5): Cis |

Each model described in Eqs. (1), (3)-(5) were fitted to the longitudinal tumour growth data from Paffenholz et al.^11^ under the four tested conditions: no treatment (vehicle), treatment with cisplatin (Cis), treatment with ICB (ICB), and treatment with cisplatin and ICB (Cis+ICB). To do so, we minimized the residual sum of squares (RSS) given by:

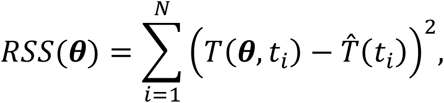

where *T*(*θ, t*_*i*_) is the model-predicted tumour volume at observation *t*_*i*_ given the parameter set *θ* and 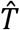 (*t*_*i*_) is the observed tumour volume at *t*_*i*_. Parameters were fitted sequentially as model complexity increased. We fitted *r* and *K* to the MP and MPB1 murine data in all four treatment scenarios using Eq. (1). We then fixed *r* and *K* values to their estimates from the vehicle data and fit Eq. (3) to the cisplatin treatment data to estimate *k*_2_.

To estimate immune cell related parameters in Eq. (4), we used the flow cytometry data of the percentage of T cells relative to CD45+ cells, the proportion of CD8+ cells, and the proportion of NK cells relative to CD45+ cells for both MP and MPB1 mice from the vehicle data, respectively, from Paffenholz et al.^11^ For this, we made several assumptions, namely that flow cytometry analyses measured 3 × 10^6^ total cells and was performed 10 days after treatment initiation, corresponding to the time of the last tumour volume measurement. We further assumed that 20-60% of the total cells analyzed were CD45+ cells^31^, to convert NK cell proportions (reported in % of CD45+ cells) to cell counts. We performed the same conversion for the CD8+ T cells (also reported in % of CD45+ cells), with the additional assumption that 20-40% of T cells are CD8+ T cells^32^ (see additional details in the Supplementary Information). We fixed the carrying capacity *K* at the value estimated by fitting Eq. (1) to the vehicle data but re-estimated the growth rate *r*, as it represents an effective net growth rate that accounts for both tumour cell production and loss in Eq. (1). With the introduction of explicit mortality terms in the tumour-immune model, *r* had to be re-estimated in Eq. (4). Parameters *p* and *q* were estimated in Eq. (4) using the vehicle data.

In the tumour-immune and cisplatin PK/PD model of Eq. (5), the immune-recruitment rates, *r*_*C,N*_ and *r*_*C,E*_ for NK and CD8+ T cells, respectively, were estimated using the cisplatin treatment-response data, while the cisplatin-induced tumour killing rate, *k*_2_, was fixed at the value estimated from the cisplatin PK/PD model in Eq. (3). Lastly, to capture the effects of ICB treatment, we re-estimated *q* in Eq. (4) using the tumour treatment-response data.

### Practical identifiability and confidence intervals

Practical identifiability analysis was performed for each model by calculating the univariate profile likelihood curves^33^ of each parameter. Let *θ* be the vector of estimated parameters and *θ*^*best*^ be the best fit estimate that minimizes the −2 log-likelihood (−2LL) given by

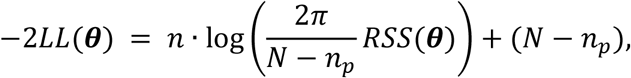

where *N* is the number of observations, *n*_*p*_ is the number of parameters to estimate in the candidate model, and *RSS(θ*) is the residual sum of squares defined above. A profile likelihood curve, *PL*_*j*_(*p*), is the −2LL minimized by all other parameters 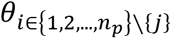, given the fixed value *p* of parameter *θ*_*j*_. Thus, *PL*_*j*_(*p*) is given by

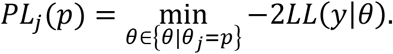

To evaluate the profile likelihood curve, for each parameter 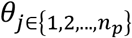, we tested 501 evenly spaced points *p* between the interval 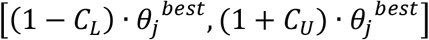. The values for the multiplicative lower and upper bound coefficients, *C*_*L*_ and *C*_*U*_, respectively, for each parameter were chosen depending on the width of their confidence intervals (see **Table S9** in the Supplementary Information).

The (1 − *α*)-level confidence interval about 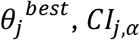, is given by the set of values of *p* for which *PL*_*j*_(*p*) is equal or less than the −2LL at *θ*^*best*^ added with the chi-square critical value 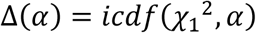, i.e.,

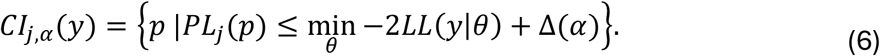

Here, we used a level of significance of *α* = 0.05.

### Model selection

To analyse parsimony and select the model most representative of the data, we calculated the corrected Akaike information criterion (AICc):

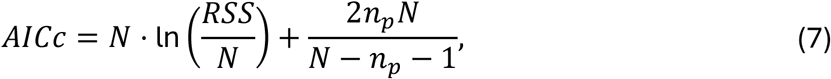

and the Bayesian information criterion (BIC):

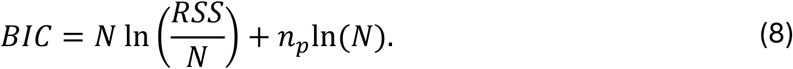

for each candidate model (Eqs. (1), (3)-(5)). To interpret the values of the criteria when comparing plausible models, we used the difference in AICc and BIC, i.e.,

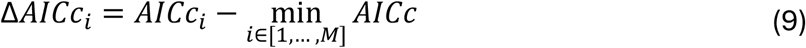

and

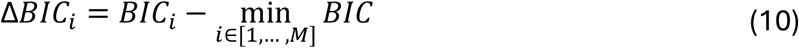

for the *M* different models. The preferred model is the one with the minimum AICc or BIC value, i.e., with a ΔAICc or ΔBIC equal to zero.

### Computational implementation

Model parameters were estimated by bound-constrained nonlinear least squares using LsqFit.jl and the *curve_fit* constructor in Julia^34^. Parameter uncertainty was assessed using the profile likelihood approach, as described above. The model ODEs were solved numerically using DifferentialEquations.jl and the *ODEProblem* constructor in Julia^34^. Model selection was directly calculated in Julia^34^. Figures were produced using the *PlotlyJS* visualization library. The code used to reproduce the analyses and figures is available on GitHub (https://github.com/Craig-Lab/MP-MPB1-mechanistic-model).

## Results

### Both MP and MPB1 tumours display saturating growth dynamics

We first investigated which growth model in Eq. (1) best described untreated tumour expansion in both MP and MPB1 mice. To do so, we fitted each growth model to the vehicle data and ranked them according to their *Δ*AICc and *Δ*BIC values (**Table S1**). Both logistic and Gompertz models closely captured the experimental tumour growth dynamics (**Figure S1**), implying that the two tumour types display saturating growth dynamics (**Figure 2**). Model selection favoured the logistic model for MP mice and the Gompertz model for MPB1 mice (see **Table 4**), albeit with small differences in both cases. Both logistic and Gompertzian growth laws are special cases of the generalized logistic framework. Since the logistic growth model has a simpler mathematical structure, we selected the logistic growth function to model tumour growth (see SI for more details). The use of a common growth model was done to facilitate direct comparison of parameter estimates between MP and MPB1 systems in the next stages of model development, where additional biological mechanisms were incorporated.

**Table 4.** Model selection results for simple tumour growth models. Calculated ΔBIC (Eq. (10)) values for the three tested growth models of Eq. (1). See **Tables S1-S4** in the Supplementary Information for full model selection results including ΔAICc values.

| Model | $\Delta BIC$ | |
| --- | --- | --- |
|  | *MP* | *MPB1* |
| **Exponential** | 23.3 | 15.1 |
| **Gompertz** | 4.2 | 0 |
| **Logistic** | 0 | 5.2 |

**Figure 2.**
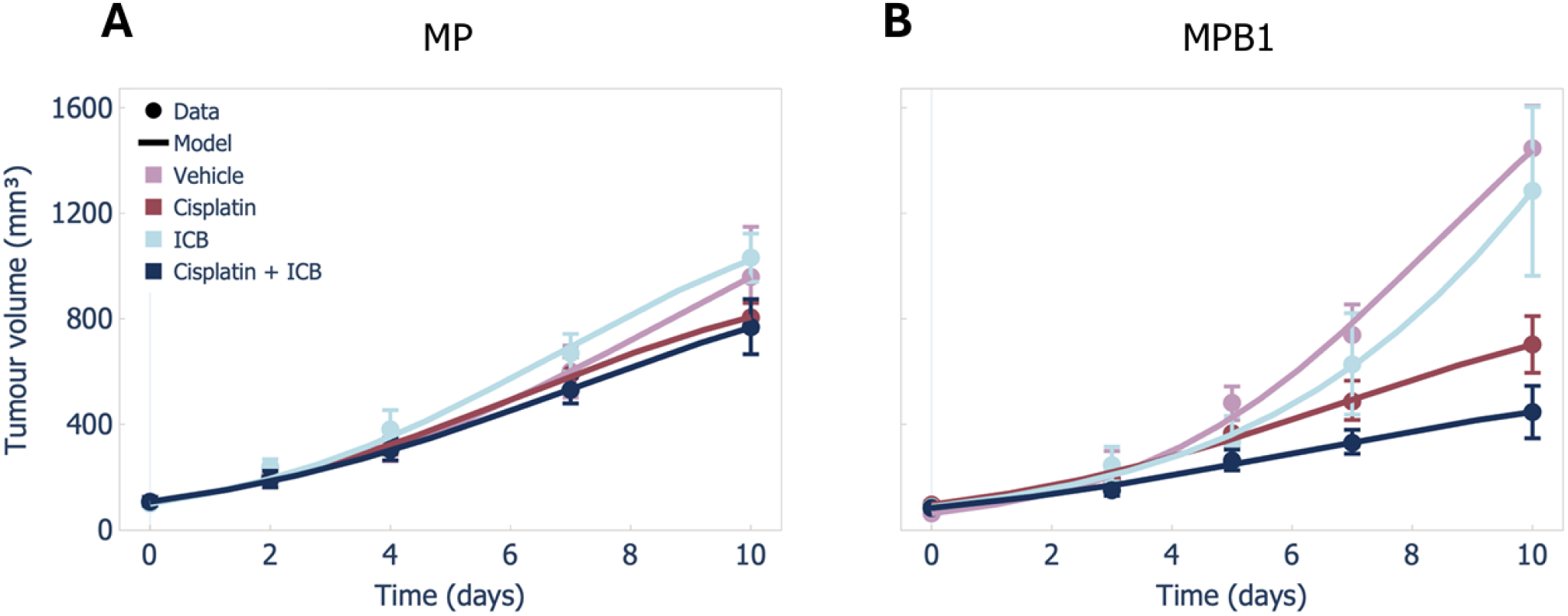
Logistic model visually captures MP and MPB1 tumour growth dynamics across treatments. Predicted tumour growth of the logistic model fit to data of vehicle (untreated), cisplatin, ICB, and cisplatin + ICB treated **A)** MP and **B)** MPB1 tumours. Despite producing visually successful fits, the correlations between the intrinsic tumour growth rate, *r*, and the tumour carrying capacity, *K*, make interpreting corresponding parameter estimation results (**Table 5**) difficult. Solid lines: model predictions; circles: mean data; error bars: standard error of the mean (SEM).

Fitting the logistic model to treated tumour volume data (i.e., cisplatin, ICB, and cisplatin + ICB) for both MP and MPB1 mice, we obtained treatment-specific estimates for the tumour growth rate, *r*, and the tumour carrying capacity, *K* (**Table 5**). All parameters were found to be practically identifiable, with the exception of *K* for MPB1 ICB-treated mice (**Figures S3-S4**). Despite providing a good visual fit to the data across all treatment regimens (see **Figure 2**), the resulting parameter values are difficult to interpret biologically because changes in *r* can be compensated by changes in *K*, producing similar tumour growth trajectories. Consequently, substantially different parameter estimates may provide similar tumour volume trajectories, as observed for the MPB1 vehicle and ICB treatment groups (**Figure 2**; **Table 5**).

**Table 5.**
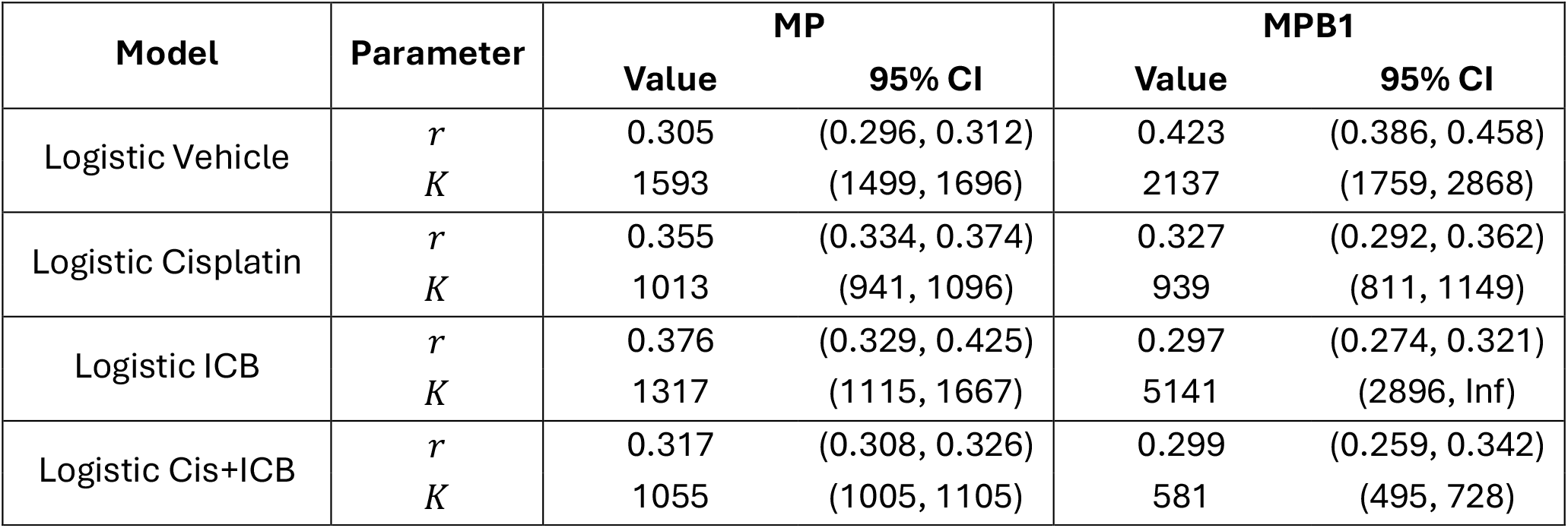
Parameter estimates of the logistic model across treatments. Values of the intrinsic tumour growth rate, *r* (in day^−1^), and tumour carrying capacity, *K* (in mm^3^), with 95% confidence intervals (CI) obtained from the profile likelihood approach for the logistic model fit to all treatments.

### Incorporating cisplatin PK/PD reveals enhanced tumour-killing effect on MPB1 tumours

To improve the interpretability of treatment effects and to discern how they may differ between MP and MPB1 mice, we extended the model to explicitly represent the actions of cisplatin on tumour growth. To quantify the effects of cisplatin treatment, we fitted the linear killing rate of cisplatin, *k*_2_, from the PK/PD model in Eqs. (2) and (3) to the same cisplatin treatment tumour volume measurements with the growth parameters, *r* and *K*, fixed to the estimates from untreated tumours (vehicle data; **Table 5**). All other parameters were set to the values reported in **Table 1 and 2**. Cisplatin administration was simulated as a bolus injection on days 0 and 7, per the experimental protocol in Paffenholz et al.^11^

In both groups, the cisplatin PK/PD model effectively captured the observed tumour growth dynamics (**Figure 3**). Notably, the model revealed differences in the estimated cisplatin PD effects between the two mouse models, particularly following the second dose at day 7, with a pronounced nonlinear response in MPB1 mice. These differences were reflected in the estimated values of parameter *k*_2_. Whereas *k*_2_ was estimated as 8.07 L*⋅*mg^−1^day^−1^ (SE = 2.63) in MP mice, it was nearly 8-times larger (63.7 L*⋅*mg^−1^day^−1^, SE = 4.56) in MPB1 mice. Thus, incorporating the cisplatin PK/PD model, despite the increase in model complexity, quantified an approximately 8-fold greater cisplatin tumour-killing effect in MPB1 compared with MP tumours.

**Figure 3.**
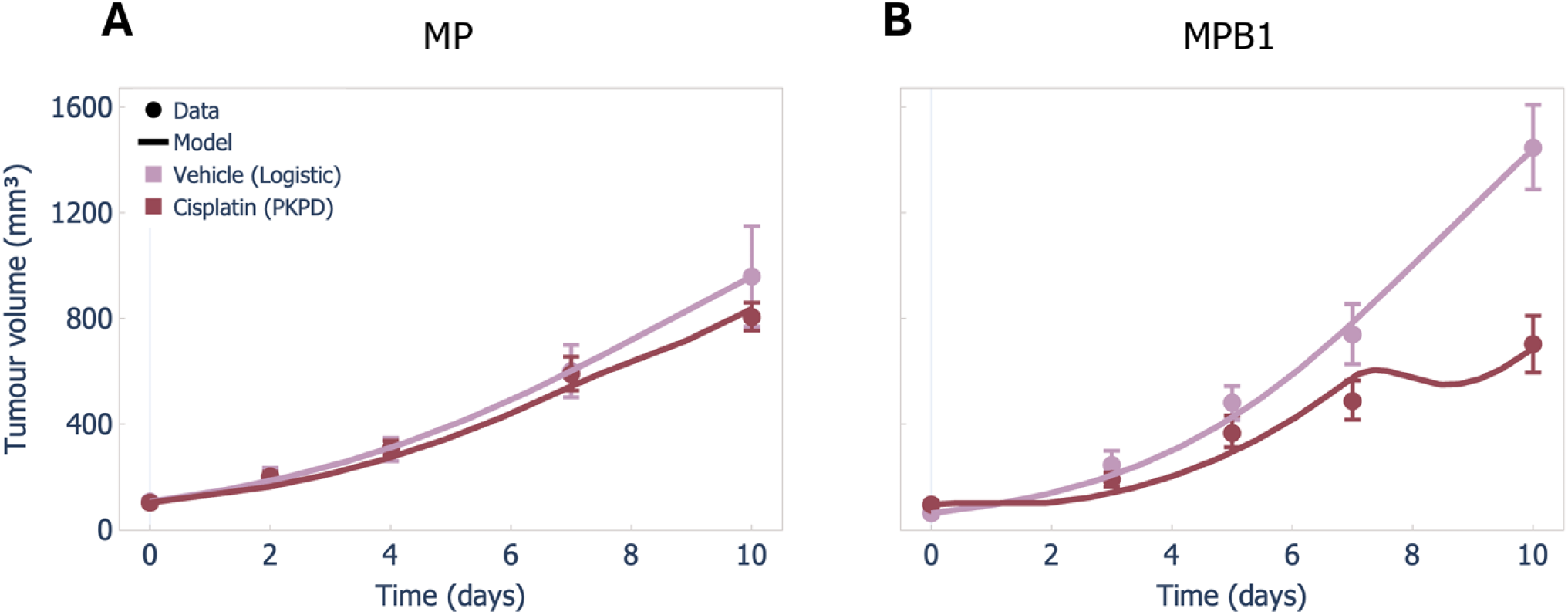
Integrating cisplatin tumour growth and cisplatin PK/PD refines biological meaning. Cisplatin PK/PD model and cisplatin-treated tumour volumes (red) compared with the logistic model and vehicle-treated tumours (pink), for **A)** MP tumours and **B)** MPB1 tumours. Solid lines: model predictions; circles: mean data; error bars: standard error of the mean (SEM).

**Figure 4.**
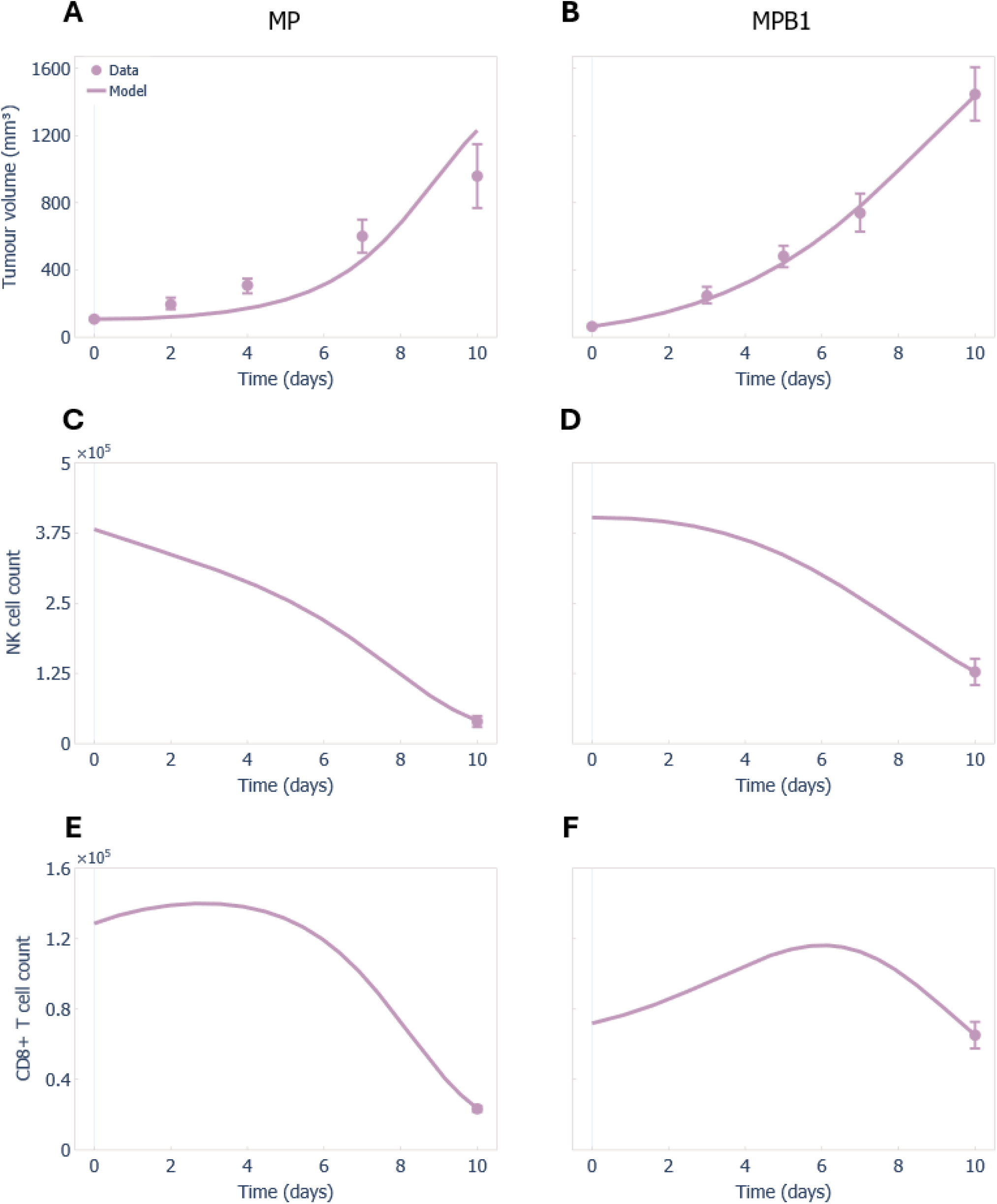
Tumour-immune extended model predicts immune cell trajectories during tumour growth. Predicted tumour growth and immune cell levels from the tumour-immune extended model (Eq. (4)) fitted to vehicle (untreated) tumour volume measurements (**A-B**), NK cell counts (**C-D**), and CD8+ T cell counts (**E-F**) in MP (**A, C, E**) and MPB1 (**B, D, F**) tumours. Solid lines: model predictions; circles: mean data; error bars: standard error of the mean (SEM).

We next calculated and compared the ΔAICc and ΔBIC values for the logistic model and the cisplatin PK/PD-integrated model fitted to cisplatin-treated MP and MPB1 tumour volume measurements. Both information criteria favoured the logistic growth model with treatment-specific parameters, *r* and *K*, over the cisplatin PK/PD model for both tumour types (**Table S2**). These results suggest that, given the available data, the logistic growth model provides a more parsimonious description of tumour growth, whereas integrating cisplatin PK/PD provides greater mechanistic insight by quantifying the anticancer effect of cisplatin for each tumour type.

### Exploring tumour-immune interactions: towards a mechanistic model of ICB treatment

Paffenholz et al.^11^ reported differences in NK and CD8+ T cell recruitment within the tumour microenvironment under vehicle and cisplatin treatment, with distinct patterns observed between MP and MPB1 mice. To construct a mechanistic model of ICB treatment, we explored how tumour-immune environment affects treatment responses by incorporating NK and CD8+ T cell interactions to our model (Eq. (4)). Because initial immune cell states were not reported in Paffenholz et al.^11^, we estimated the time between tumour and treatment initiation per the logistic growth model (Eq. (1)) and inferred NK and CD8+ T cell counts at treatment initiation (see Supplementary Information, Figure S2). We then jointly estimated the tumour growth rate, *r*, and the rates of NK and CD8+ T cell inactivation, *p* and *q*, respectively, by fitting this model to tumour volume dynamics and NK and CD8+ T cell count data from both MP and MPB1 mice. All other parameter values were fixed to previously reported values (**Table 1 and 2**), except for the tumour carrying capacity, *K*, which was fixed at the values previously estimated from the vehicle data using Eq. (1) (see **Table 5**).

The extended tumour-immune model successfully recapitulated observed tumour growth dynamics and immune cell counts (**Figure 4**). The estimated growth rates, *r*, were found to be higher (**Table 6**) than those obtained from the logistic growth model (**Table 4**). This is to be expected as Eq. (4) explicitly describes tumour cell killing/removal by immune cells, whereas these are absent in the simple logistic growth model in Eq. (1). This results in compensation in the net tumour growth rate *r* = *b* − *d*, where *b* represents the tumour production rate and *d* the death rate. Tumours in MPB1 mice nevertheless showed consistently higher growth rates than in MP mice, as before. The estimated immune cell inactivation rates, *p* and *q*, were also higher in MP mice than in MPB1 mice tumours (**Table 6**). As these parameters govern the inactivation of NK and CD8+ T cells within the model, the estimates are consistent with the lower immune cell counts observed in the MP mice tumour microenvironment. All parameters were found to be practically identifiable (**Figure S5**).

**Table 6.**
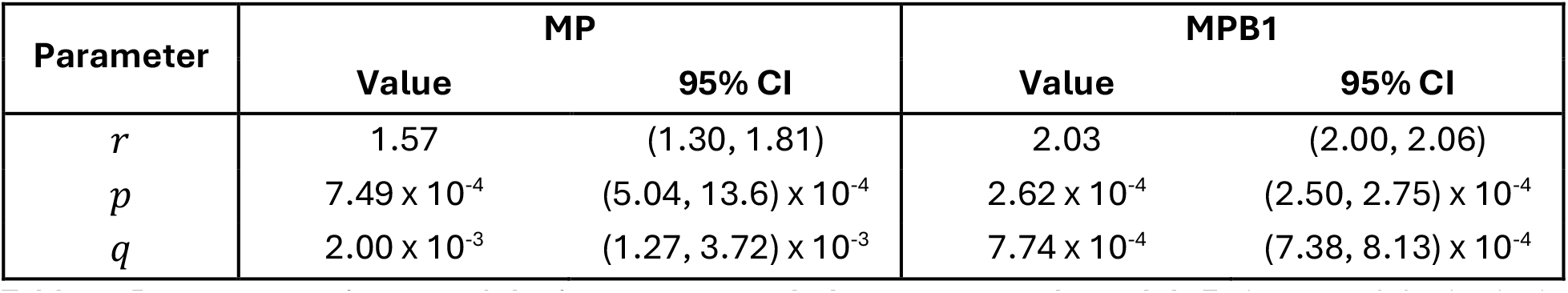
Parameter estimates of the immune-extended tumour growth model. Estimates of the intrinsic tumour growth rate, *r* (day^−1^), and the NK cell and CD8+ T cell inactivation rates due to tumour interaction, *p* and *q*, respectively (mm^−3^ day^−1^). Values are reported with 95% confidence intervals (CI) obtained from the profile likelihood approach.

Unfortunately, model selection criteria cannot be used to compare the immune-extended model with the logistic growth model, as the two models were fitted to different datasets with different numbers of observations. Nevertheless, the tumour-immune extended model (Eq. (4)) provided a fit to MPB1 data comparable to that of the logistic model (RSS = 3.86 × 10^3^ vs. 6.80 × 10^3^, respectively), although its fit to MP tumour volume data was substantially poorer (RSS = 1.24 × 10^5^ vs. 88.4, respectively). Importantly, the extended model predicted immune-cell differences consistent with those reported between MP and MPB1 mice, while providing a framework for modelling immune-mediated cisplatin and ICB treatment.

### The full tumour-immune model integrating cisplatin PK/PD successfully captures cisplatin treatment effects

Next, to investigate the role of NK and CD8+ T cells during cisplatin treatment, we fit the integrated PK/PD-immune model in Eq. (5) to tumour volume, NK cell count, and CD8+ T cell count data from cisplatin-treated MP and MPB1 mice^11^. For this, we again fixed most parameters to the values reported in **Table 1 and 2** and set the carrying capacity, *K*, to the values previously estimated by fitting the logistic model to vehicle data (**Table 5**). Parameters *r, p*, and *q* were fixed to the values estimated using the immune-extended model under vehicle treatment described in the previous section (**Table 6**).

In de Pillis et al.^35^, cisplatin-induced cytotoxicity was assumed to act uniformly on tumour and immune cells through shared mortality terms (*k*_2_) applied to NK and CD8+ T cell. However, Paffenholz et al.^11^ reported increased immune cell infiltration following cisplatin treatment in certain tumour types, suggesting a potential drug-induced enhancement of immune recruitment in the tumour microenvironment. To account for this effect, we extended the model by introducing cisplatin-dependent recruitment terms for NK cells and CD8+ T cells, *r*_*C,N*_ and *r*_*C,E*_ respectively (Eq. (5)). In this case, the net recruitment rates are therefore given by *r*_*C,N*_ − *k*_2_ and *r*_*C,E*_ − *k*_2_, allowing for either net depletion or net recruitment depending on parameter values.

We took two approaches to the integrated cisplatin PK/PD-immune model (**Figure 5**). First, we assumed there was no effect of cisplatin on the immune cells (i.e., we set *r*_*C,N*_ = *r*_*C,E*_ = *k*_2_) and simulated Eq. (5) using the value of *k*_2_ previously estimated from cisplatin PK/PD model (*k*_2_ = 8.07 L*⋅*mg^−1^day^−1^ for MP tumours, *k*_2_ = 63.7 L*⋅*mg^−1^day^−1^ for MPB1 tumours; **Table S5**). This way, cisplatin effects were only modelled as affecting the mortality of the tumour cells. With the same fixed *k*_2_ values, we also estimated the rate of NK cell recruitment, *r*_*C,N*_, and the rate of CD8+ T cell recruitment, *r*_*C,E*_, from the tumour volume and immune cell count data^11^, thereby incorporating the cisplatin-mediated modulation of immune cell recruitment in the model estimates.

**Figure 5.**
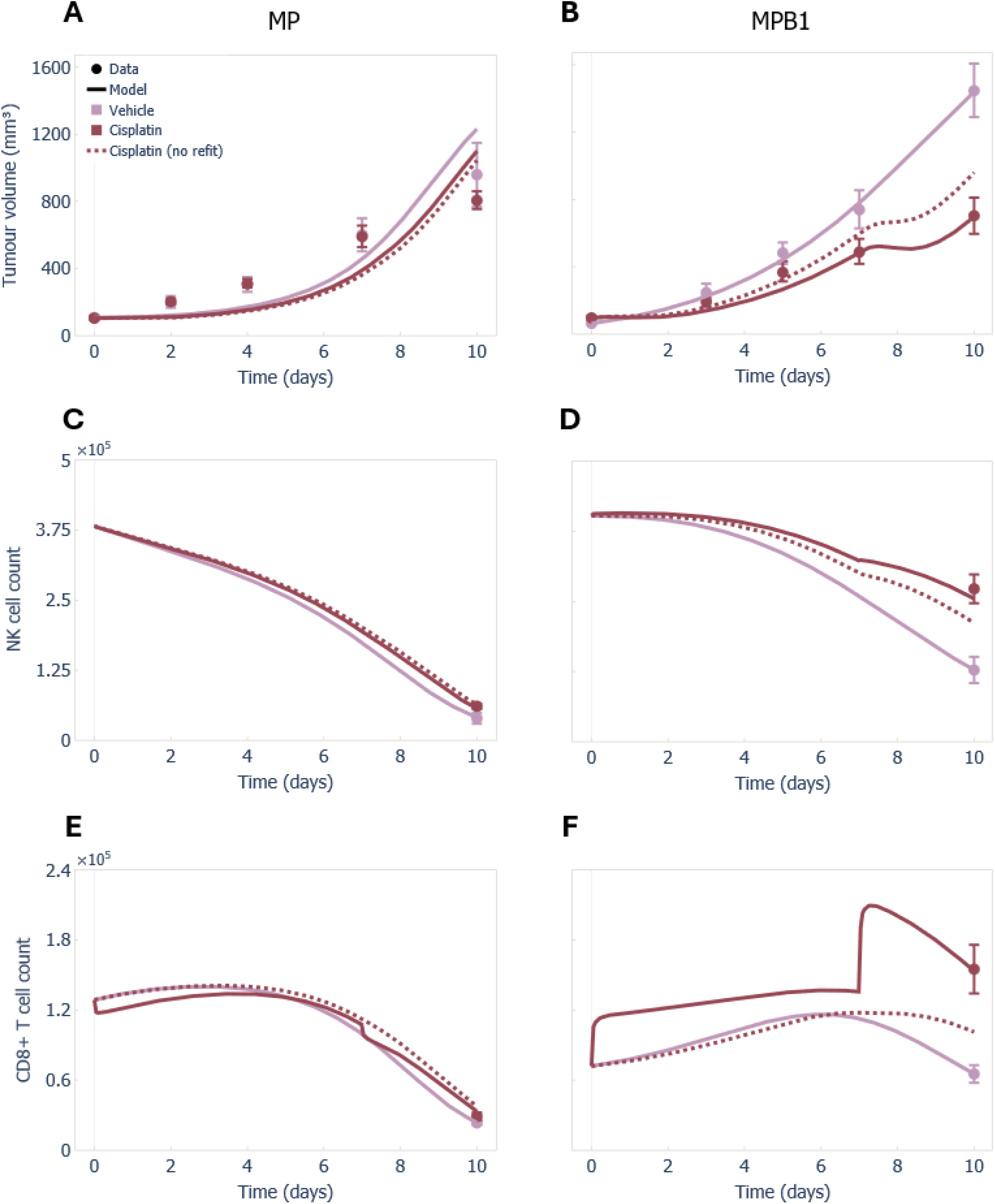
Predicted cisplatin-induced immune recruitment affects tumour and immune cell dynamics in MPB1 but not MP tumours. Combined tumour-immune and cisplatin PK/PD model (Eq. (5)) predictions without additional fitting (red dotted lines) and with fitting of the cisplatin-induced immune recruitment terms (red solid lines), compared with the vehicle model fits of Eq. (4) (pink), for tumour volume measurements (**A-B**), NK cell counts (**C-D**), and CD8+ T cell counts (**E-F**) in MP (**A, C, E**) and MPB1 (**B, D, F**) tumours. Solid and dotted lines: model predictions; circles: mean data; error bars: standard error of the mean (SEM).

The estimated cisplatin-induced immune recruitment parameters indicated a marked increase in immune cell recruitment in MPB1 tumours versus close to zero in MP tumours (**Table 7**). Further, parameters were found to be practically identifiable only for the MPB1 tumours (**Figure S6**). Consistent with this, the 95% confidence intervals were narrow for the parameters in MPB1 tumours (**Table 7**), whereas those for the MP tumour were considerably wider, with only an upper confidence bound that could be estimated, reflecting their practical unidentifiability.

**Table 7.** Parameter estimates of the immune-extended tumour growth and cisplatin PK/PD model. Estimates of the NK cell and CD8+ T cell recruitment rates due to cisplatin, *r*_*C,N*_ and *r*_*C,E*_, respectively (L*⋅m*g^−1^day^−1^). Values are reported with 95% confidence intervals (CI) obtained from the profile likelihood approach, with no lower bounds found for MP estimates.

| Parameter | MP |  | MPB1 |  |
| --- | --- | --- | --- | --- |
|  | Value | 95% CI | Value | 95% CI |
| $r_{C,N}$ | 7.95 | (-inf, 9.24) | 64.4 | (62.4, 66.2) |
| $r_{C,E}$ | 0 | (-inf, 81.2) | 97.9 | (77.4, 113.7) |

The estimated net effect of cisplatin on NK cells (i.e., *r*_*C,N*_ − *k*_2_) was close to zero in both MP and MPB1 tumours, with point estimates of −0.1 and 0.7 L*⋅*mg^−1^day^−1^, respectively. The confidence intervals for *r*_*C,N*_ included values corresponding to both positive and negative net effects, preventing a definitive conclusion regarding cisplatin-mediated NK cell modulation in both tumour types. The estimated net effect of cisplatin on CD8+ T cells (*r*_*C,E*_ − *k*_2_) was positive for MPB1 tumours, despite the higher estimated value of *k*_2_, with a net recruitment of 34.2 L*⋅*mg^−1^day^−1^. For MP tumours, no reliable inference could be made, but the model fits with the best fit value *r*_*C,E*_ = 0 (**Figure 5**) were consistent with a null or very limited cisplatin-induced recruitment of T cells. These results suggest that cisplatin-induced modulation of the tumour immune microenvironment is more pronounced in MPB1 than in MP tumours. As before, model selection criteria cannot be used to directly compare the cisplatin PK/PD-immune model and the cisplatin PK/PD or logistic growth models, as they were fitted to different sets of observations.

### A fully mechanistic model of ICB without and with cisplatin treatment

To investigate the mechanisms of immune checkpoint blockade, we fitted the extended tumour-immune model in Eq. (4) to tumour volume data from ICB-treated mice by re-estimating the tumour-mediated inactivation of CD8+ T cells, *q*. For this, we fixed the carrying capacity, *K*, to its estimate from the logistic growth model (**Table 5**), while the growth rate, *r*, and the rate of NK cell inactivation by the tumour, *p*, were fixed to estimates obtained from the extended tumour-immune-fit to the vehicle treatment (**Table 6**). All other parameters were set to the values reported in **Table 1**. In MP tumours treated with ICB, the value of *q* was comparable to the vehicle tumour (2.14 × 10^−3^ mm^−3^day^−1^ vs. 2.00 × 10^−3^ mm^−3^day^−1^, respectively; **Table S5**). In contrast, *q* was significantly reduced in MPB1 tumours treated with ICB compared to the untreated MPB1 tumours (1.41 × 10^−4^ mm^−3^ day^−1^ vs. 7.74 × 10^−4^ mm^−3^day^−1^, respectively; **Table S5**). These results suggest that ICB treatment has little effect in the tumour-immune microenvironment of MP tumours, whereas it effectively reduces CD8+ T cells inactivation in MPB1 tumours (**Figure 6**). Comparing the ΔAICc and ΔBIC values (see **Table S3**), the logistic growth models with ICB treatment-specific parameters *r* and *K* (**Table 5**; **Figure 2**) were also favoured over the tumour-immune model with ICB-specific parameters (**Figure 6**).

**Figure 6.**
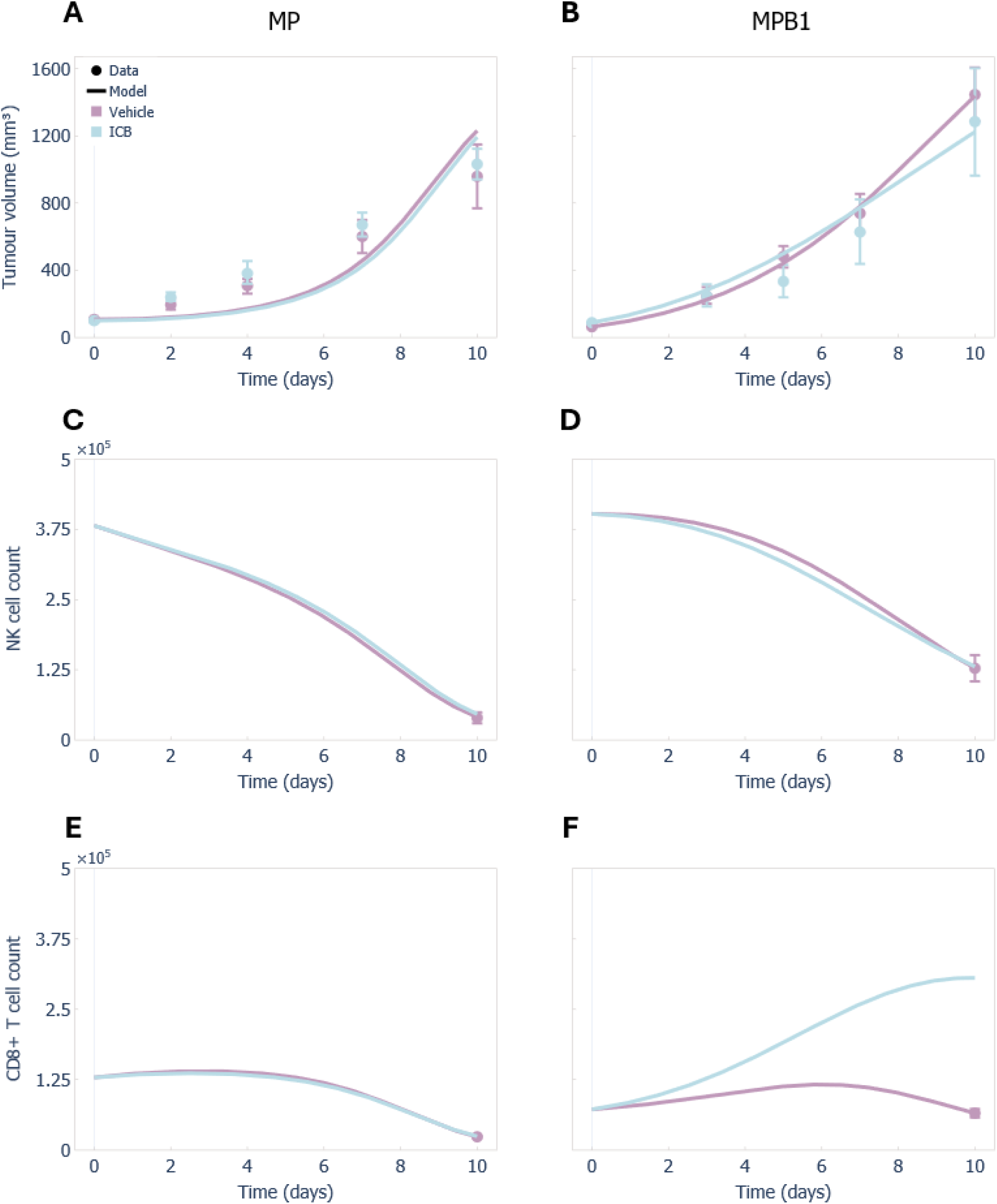
Model predicts reduced tumour-mediated T cell inactivation and increased T cell counts under ICB in MPB1 but not in MP tumours. Predicted tumour growth and immune cell levels from the tumour-immune extended model (Eq. (4)) fitted to vehicle (pink) and ICB-treated (light blue) tumour volume data, with only the tumour-mediated T cell inactivation rate re-estimated for ICB-treated tumours. Model predictions are shown for tumour volume (**A-B**), NK cell counts (**C-D**), and CD8+ T cell counts (**E-F**) in MP (**A, C, E**) and MPB1 (**B, D, F**) tumours. Solid lines: model predictions; circles: mean data; error bars: standard error of the mean (SEM). No immune cell counts were available for ICB-treated mice for model fitting.

Finally, to validate the predictive ability of the mechanistic model, we used Eq. (5), combining logistic tumour growth, cisplatin PK/PD, NK and CD8+ T cell interactions, and the ICB effects, to predict the response to combination therapy (cisplatin + ICB) observed in the experimental data^11^. Parameters estimated from previous model fits were fixed to the corresponding estimates summarized in **Table S5**. Specifically, we fixed *K* to the estimates obtained by fitting the logistic growth model (Eq. (1)) to vehicle-treated data, and *k*_2_ to the estimates obtained by fitting the cisplatin PK/PD model (Eq. (3)) to cisplatin-treated data. Parameters *r* and *p* were fixed to the estimates obtained by fitting the tumour-immune extended model (Eq. (4)) to vehicle-treated data, and *q* to the estimates obtained by fitting Eq. (4) to ICB-treated data. Parameters *r*_*C,N*_ and *r*_*C,E*_ were fixed to the estimates obtained by fitting the tumour-immune and cisplatin PK/PD model (Eq. (5)) to cisplatin-treated data. All other parameters were fixed to the values reported in **Table 1 and 2**.

The combined model incorporating cisplatin PK/PD and ICB-informed parameter estimates yielded a close match to the experimental tumour volume data for MPB1 tumours, and similar tumour growth trajectories to those observed under the other treatment conditions for MP tumours (**Figure 7**). Model selection criteria again favoured the logistic growth model over the combined model for cisplatin + ICB-treated tumour growth data, with a smaller difference between the models for MPB1 tumours (see **Table S4**).

**Figure 7.**
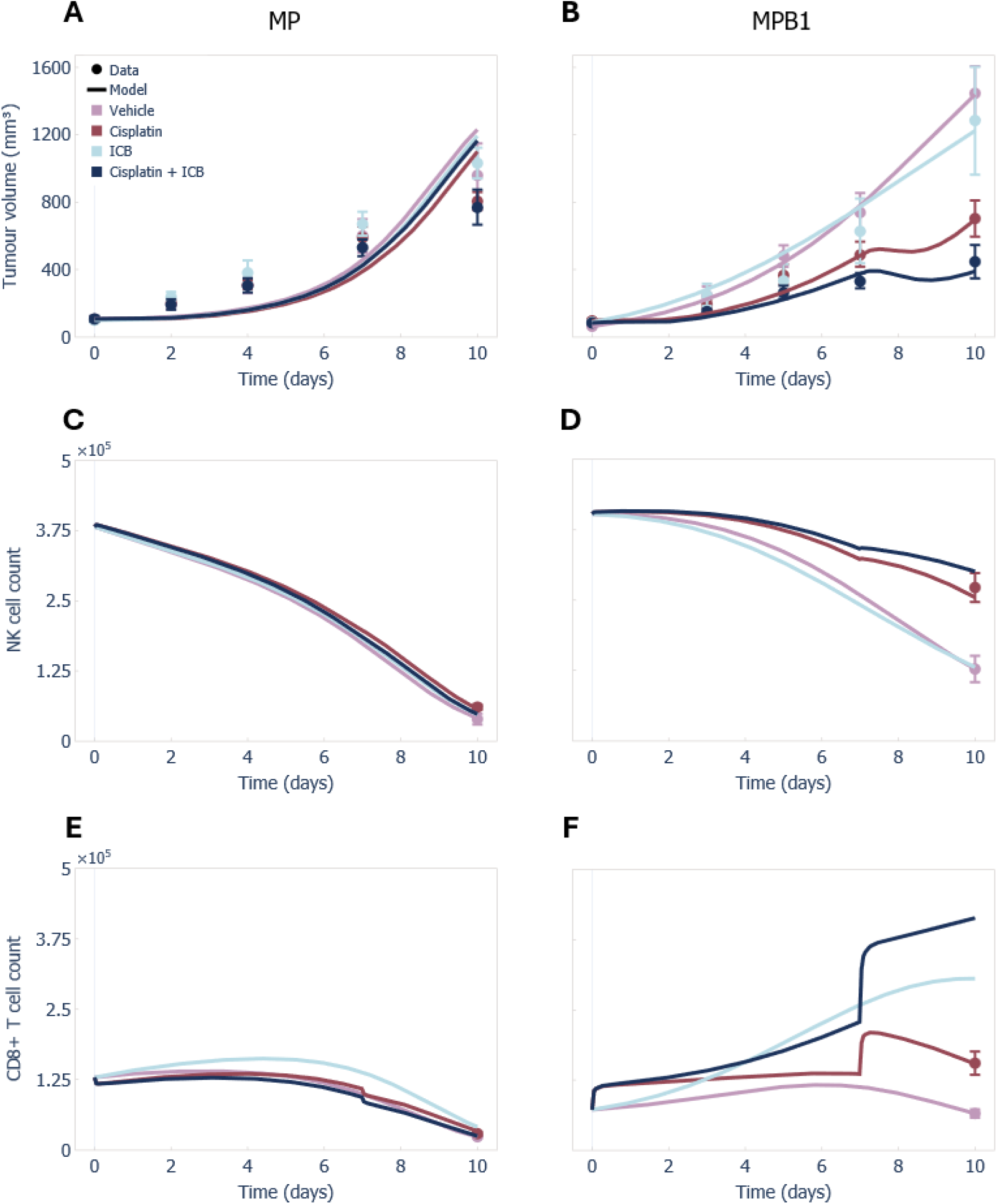
Combined mechanistic model predicts tumour response to cisplatin + ICB treatment. Model predictions for tumour growth and immune cell trajectories under vehicle (pink) and ICB (light blue) treatments using the tumour-immune extended model (Eq. (4)), and under cisplatin (red) and cisplatin + ICB (dark blue) treatments using the combined tumour-immune and cisplatin PK/PD model (Eq. (5)). Parameter values used for the predictions are summarized in **Table 1 and 2** and **Table S5**. Predictions are compared with experimental tumour volume measurements (**A-B**), NK cell counts (**C-D**), and CD8+ T cell counts (**E-F**) in MP (**A, C, E**) and MPB1 (**B, D, F**) tumours. Solid lines: model predictions; circles: mean data; error bars: standard error of the mean (SEM). No immune cell counts were available for ICB and cisplatin + ICB treatment conditions.

## Discussion

Mathematical modelling can help uncover underlying biological interactions, complementing experimental and clinical approaches. However, there is a delicate balance to be struck between using the simplest model that adequately describes the experimental data and developing more complex mechanistic mathematical models that enable predictive extrapolation. Here, we studied a hierarchy of mathematical models of increasing biological complexity to elucidate the interplay between model complexity and understanding treatment effects in two mice models of high-grade serous ovarian cancer. While simpler models adequately described tumour growth dynamics between tumour types in the absence of treatment, more mechanistic models were required to capture immune cell recruitment and to investigate treatment effects, such as those induced by immune checkpoint blockade. Overall, our results highlight how different levels of model complexity serve reciprocal purposes with different predictive capacities. Based on our results, we therefore advocate for constructing fit-for-purpose mechanistic models that answer specific biological questions of interest, rather than purely relying on statistical measures of model selection.

Model selection criteria, such as AICc and BIC, are fundamental components for comprehensive model selection. These statistical approaches favoured tumour growth models incorporating a saturation term to better represent the data. Interestingly, differences between tumour types were already evident, with the logistic growth favoured for MP tumours and the Gompertz growth for MPB1 tumours (**Table 4**). When a single model was selected to facilitate comparison between tumour types, the estimated parameters still differed substantially, with higher growth rates and carrying capacities for the MPB1 tumours (**Table 5**), suggesting that MPB1 tumours grow faster and reach larger sizes than MP tumours, a prediction that was not previously explicitly investigated^11^.

Irrespective of the modelling framework, we found that cisplatin exhibits distinct effects on MP and MPB1 tumours. Under a logistic growth model, the parameter estimates for the growth rate and carrying capacity for cisplatin-treated MPB1 tumours were considerably lower than those estimated for the vehicle-treated tumours (**Figure 2**; **Table 5**), a clear indication that MPB1 tumours are sensitive to cisplatin. Comparatively, there was a decrease in the carrying capacity but an increase in the growth rate in MP tumours (**Table 5**). However, interpreting these treatment effects is difficult because the growth rate and carrying capacity have compensatory influences on tumour dynamics. In contrast, dissecting the predicted effects using the cisplatin PK/PD model (Eqs. (2) and (3), **Figure 3**) is more straightforward. Indeed, this model predicted a relatively small cisplatin-induced mortality rate for MP tumours, indicating limited sensitivity to treatment, whereas the estimated mortality rate was approximately eightfold higher for MPB1 tumours. The PK/PD formulation therefore enabled direct comparison between vehicle- and cisplatin-treated tumours as well as between MP and MPB1 tumours. Although standard model selection criteria favoured the logistic growth model because it achieved a better fit to the data, the PK/PD model proved more informative biologically by providing directly interpretable treatment-effect parameters. This illustrates that the statistically best-fitting model is not necessarily the most useful one for addressing a biological question.

Incorporating immune cell interactions to the tumour growth model and calibrating the model resulted in the estimation of two- or three-fold higher immune cell inactivation rates due to tumour interactions in MP versus MPB1 tumours (**Table 6**). Although the qualitative difference of immune cell infiltration between tumour types was already apparent from the experimental observations^11^, this extended model provided a quantitative estimate of its magnitude. More importantly, it establishes a framework for addressing subsequent biological questions, particularly for comparing the mechanisms and effects of ICB.

HR-deficient tumours (i.e., MPB1) may be more receptive to immunotherapy than HR-proficient tumours (i.e., MP) because they tend to have higher neoantigen loads, more tumour-infiltrating lymphocytes, and higher immune-pathway gene expression. In contrast, MP tumours have a more immunosuppressive microenvironment, reducing immunotherapy efficacy^36,37^. Consistent with this biology, our extended model predicted an increase in CD8+ T cell counts in ICB-treated MPB1 tumours (**Figure 6**). Comparing the ΔAICc and ΔBIC values (**Table S3**), the logistic growth models with ICB treatment-specific *r* and *K* values were favoured over the tumour-immune model with ICB-specific parameters (with ΔBIC values between 14-17). Although statistical comparison with the logistic growth model informed by the same treatment data favoured the simpler formulation in terms of descriptive fit, the tumour-immune model provides a mechanistic interpretation of treatment response. Specifically, fitting the extended model fit to ICB treatment data revealed a marked (approximately five-fold) reduction in the tumour-induced CD8+ T cell inactivation rate in MPB1 tumours compared to vehicle data, while this parameter remained mostly unchanged in MP tumours. This indicates a measurable predicted treatment effect, despite the lack of a statistically significant difference between vehicle and ICB-treated MPB1 tumours in the experimental results^11^. In particular, the reduced CD8+ T cell inactivation suggests that ICB may affect MPB1 tumour growth by reducing tumour-induced immune suppression, providing a roadmap for future experimental investigation. Hence, the more complex mechanistic model allowed for the identification of differential changes in the tumour-induced CD8+ T cell inactivation rate between MP and MPB1 tumours under ICB treatment, offering biological insight that cannot be obtained from the simpler model. Interestingly, although our extended immune model does not explicitly describe PD-1/PD-L1 dynamics^25^, it nonetheless captured the effects of ICB, highlighting its utility when immune cell data are limited.

By incorporating immune cell dynamics from Eq. (5) into the cisplatin PK/PD model, we reproduced the immune cell counts reported by Paffenholz et al.^11^, enabling quantitative comparisons of immune cell infiltration between vehicle and cisplatin-treated MP and MPB1 tumours. Two approaches were considered using the previously estimated cisplatin PK/PD parameter *k*_2_: (i) simulation with no direct effect of cisplatin on immune cells, and (ii) estimation of the cisplatin-induced immune recruitment parameters. While tumour growth dynamics were adequately captured in both cases, immune cell dynamics in the MPB1 tumour were not reproduced when immune recruitment was omitted (**Figure 5**). In contrast, the refitted model revealed markedly different immune cell recruitment rates between cisplatin-treated MP and MPB1 tumours (**Table 7**). The estimated net effect of cisplatin on CD8+ T cells was strongly positive in MPB1 tumours. Although the corresponding parameter was not practically identifiable in MP tumours, the fitted model was consistent with the knowledge that little or no cisplatin-induced recruitment of CD8+ T cells occurs in MP tumours. Overall, these results suggest that the superior response of MPB1 tumours arise from a combination of enhanced direct cytotoxicity and treatment-induced immune recruitment, whereas both mechanisms are attenuated in MP tumours, highlighting the importance of explicitly modelling immune dynamics to capture this treatment response.

Simulating the extended cisplatin PK/PD and immune response model under cisplatin and ICB combination treatment further supports its validity, as it reproduced the observed behaviour in MPB1 tumours without requiring additional parameter re-fitting (**Figure 7**). While the model fit to MP tumours is less accurate, the relatively small differences observed experimentally in MP tumour growth between treatment conditions suggest limited treatment-driven changes in growth dynamics, consistent with the model predictions for the combination therapy. A comparison between the simpler logistic model and the mechanistic tumour– immune formulation (**Figure 2**; **Figure 4**) further highlights the differences in model structure and interpretation. Although the simpler models fit the MP tumour growth data very well, the more sophisticated mechanistic models provide greater biological interpretability, as discussed above.

MPB1 tumours recapitulate key biological and clinical features of human HR-deficient tumours^11^, which account for more than a third of ovarian cancer cases^38^. Across all modelling frameworks, our results consistently identified biological differences between MP and MPB1 tumours. Even in the absence of treatment, MPB1 tumours were predicted to have higher growth rates and carrying capacities than MP tumours. Incorporating immune cell interactions further suggested that MP tumours exert stronger tumour-mediated immune suppression, with two-to three-fold higher immune cell inactivation rates than MPB1 tumours. Under ICB treatment, this tumour-induced CD8+ T cell inactivation remained essentially unchanged in MP tumours but decreased approximately five-fold in MPB1 tumours, indicating that ICB preferentially relieves immune suppression in the HR-deficient model. Finally, the cisplatin PK/PD model estimated an approximately eightfold greater direct cisplatin-induced tumour cell mortality in MPB1 tumours than in MP tumours and predicted a strong cisplatin-induced recruitment of CD8+ T cells in MPB1, whereas the net effect on CD8+ T cells was suspected to be negative in MP tumours. Together, these findings suggest that the enhanced therapeutic response of MPB1 tumours arises from both increased intrinsic sensitivity to cisplatin and a more favourable anti-tumour immune response, consistent with the known biology of HR-deficient ovarian cancer.

Care should be taken in interpreting our results. Biologically speaking, the assumption that cisplatin targets only cycling tumour cells, *T*_1_, in the full model described in Eq. (5) discounts known effects of cytotoxic chemotherapy on immune cell populations. In terms of modelling approach, we intentionally used a hierarchical model construction to illustrate how statistical model selection techniques and model complexity interact. However, this required fixing many model parameters to values from different tumour and animal types, introducing uncertainty. The ideal scenario where we would have sufficient data to fully parameterize our models is possible^39,40^ but rare. A further limitation is that only a subset of model parameters was allowed to differ between the MP and MPB1 tumours, while the remaining parameters were assumed to be common across tumour types. The choice of which parameters were tumour-specific was informed by the biological interpretation of the experimental results provided by Paffenholz et al.^11^ Restricting the number of tumour-specific parameters was necessary to ensure that the available data was sufficient for reliable parameter estimation and to avoid an unidentifiable model. Consequently, the estimated differences between tumour types should be interpreted as the effects of the selected biological mechanisms represented by the fitted parameters, rather than as a complete description of all biological differences. If additional tumour-specific mechanisms also differ between MP and MPB1, some of their effects may be absorbed into the estimated parameter values, potentially influencing their biological interpretation.

Although most fitted parameters were practically identifiable, NK cell and CD8+ T cell cisplatin-induced recruitment in the MP tumour was poorly identifiable. This lack of practical identifiability indicates that the available data does not provide sufficient evidence to estimate a non-zero recruitment effect in this tumour type, supporting the selection of the simpler model. In contrast, the corresponding parameter was practically identifiable in the

MPB1 tumour, suggesting that the data support the inclusion of this mechanism only for the HR-deficient tumour. This demonstrates the utility of the profile likelihood approach in model selection, as it allows the identifiability of individual mechanisms to be assessed and provides a quantitative basis for determining whether additional model complexity is supported by the data.

More broadly, our results highlight that model selection should be guided by both statistical performance and the scientific question being addressed. In all cases, the simpler logistic growth model was favoured over the more mechanistic tumour-immune model. While this choice is appropriate if the objective is solely to describe tumour growth, the more mechanistic models enabled interpretation of biological mechanisms that are inaccessible to the simpler growth models, including immune cell recruitment, tumour-mediated immune suppression, and treatment-induced immune responses. Our results demonstrate that the extended models could yield substantially greater biological insight, even when improvements in goodness-of-fit were limited. Overall, our study highlights that model selection should balance parsimony with the ability of a model to address the underlying biological questions, rather than relying exclusively on statistical criteria. This is particularly critical as models are increasingly relied upon for regulatory^41^ and clinical decision making^42^, highlighting the importance of anchoring models to fundamental biological mechanisms to improve the reliability of their predictions.

## Code availability statement

The code to reproduce the analyses and figures in this study is available on GitHub (https://github.com/Craig-Lab/MP-MPB1-mechanistic-model).

## Acknowledgements

The authors would like to thank Gabriel Côté for his input at the beginning of the project.

## Author contributions

PL: Conceptualization, methodology, software, validation, formal analysis, investigation, writing, visualization, project administration.

MB: Conceptualization, methodology, software, validation, formal analysis, investigation, writing, visualization.

TE: Conceptualization, methodology, software, validation, data curation, writing, visualization.

FB: Conceptualization, methodology, software, validation, formal analysis, investigation, writing, visualization.

MC: Conceptualization, formal analysis, writing, visualization, supervision, project administration, funding acquisition.

