## SupplementaryInformation for "The trade-off between parsimony and model complexity for understanding biomedical mechanisms from mathematical models" (from another version of this preprint: version 1, 2026-09-01)

*Model Selection*

The **Tables S1–S4** summarize the model selection results corresponding to each stage of model development presented in the main text. Each table holds results for MP and MPB1 consisting of the model; number of parameters, $N$; corrected Akaike information criterion, AICc; difference in AICc, ΔAICc; Bayesian information criterion, BIC; difference in BIC, ΔBIC; and residual sum of squares, RSS.

| Tumour | Model | N | AICc | ΔAICc | BIC | ΔBIC | RSS |
|---|---|---|---|---|---|---|---|
| MP | Exponential | 1 | 42.55685 | 18.19546 | 40.83295 | 23.25269 | 12760.9 |
| | Logistic | 2 | 24.36139 | 0 | 17.58026 | 0 | 88.38612 |
| | Gompertz | 2 | 28.55255 | 4.191162 | 21.77142 | 4.191162 | 204.3731 |
| MPB1 | Exponential | 1 | 50.85982 | 10.02664 | 49.13592 | 15.08387 | 67153.4 |
| | Logistic | 2 | 46.07852 | 5.245348 | 39.2974 | 5.245348 | 6803.161 |
| | Gompertz | 2 | 40.83318 | 0 | 34.05205 | 0 | 2382.899 |

**Table S1. Model selection of tumour growth.** Selection criteria results for tumour growth models of Eq. (1) in the main text.

| Tumour | Model | N | AICc | ΔAICc | BIC | ΔBIC | RSS |
|---|---|---|---|---|---|---|---|
| MP | Logistic Cisplatin | 2 | 32.15919 | 0 | 25.37807 | 0 | 420.4295 |
| | Cisplatin PK/PD | 1 | 38.50636 | 6.347165 | 36.78246 | 11.40439 | 5676.233 |
| MPB1 | Logistic Cisplatin | 2 | 37.40934 | 0 | 30.62822 | 0 | 1201.476 |
| | Cisplatin PK/PD | 1 | 43.8393 | 6.429964 | 42.11541 | 11.48719 | 16492.02 |

**Table S2. Model selection of logistic growth and cisplatin PK/PD models.** Selection criteria results for the cisplatin-treated logistic growth model of Eq. (1) and cisplatin PK/PD model of Eqs. (2)-(3) in the main text.

| Tumour | Model | N | AICc | ΔAICc | BIC | ΔBIC | RSS |
|---|---|---|---|---|---|---|---|
| MP | Logistic ICB | 2 | 42.98889 | 0 | 36.20777 | 0 | 3667.318 |
| | Immune-extended | 1 | 54.93819 | 11.94930 | 53.21430 | 17.00653 | 151813.8 |
| MPB1 | Logistic ICB | 2 | 40.84104 | 0 | 34.05992 | 0 | 2386.649 |
| | Immune-extended | 1 | 49.68782 | 8.846785 | 47.96393 | 13.90401 | 53121.46 |

**Table S3. Model selection of logistic growth and immune-extended ICB treatment models.** Selection criteria results for the ICB-treated logistic growth model of Eq. (1) and the ICB-treated immune-extended tumour growth model of Eq. (4) in the main text.

| Tumour | Model | N | AICc | ΔAICc | BIC | ΔBIC | RSS |
|---|---|---|---|---|---|---|---|
| MP | Logistic Cisplatin + ICB | 2 | 23.74388 | 0 | 16.96275 | 0 | 78.11744 |
| | Immune-extended | 0 | 52.88361 | 29.13973 | 52.88361 | 35.92086 | 196056.8 |
| MPB1 | Logistic Cisplatin + ICB | 2 | 33.77222 | 0 | 26.9911 | 0 | 580.4955 |
| | Immune-extended | 0 | 37.02221 | 3.24999 | 37.02221 | 10.03111 | 8216.340 |

**Table S4. Model selection of logistic growth model and immune-extended cisplatin and ICB combination treatment models.** Selection criteria for the cisplatin and ICB-treated logistic growth model of Eq. (1) and the cisplatin PK/PD and immune-extended tumour growth of Eq. (5) in the main text, with parameters informed from the previous cisplatin PK/PD and ICB model fits.

*Estimated parameter values*

The **Table S5** summarizes the estimated parameter values and standard error from the fitting procedure in each stage of model development.

| Model | Fitted parameter description and units | MP | | MPB1 | |
|---|---|---|---|---|---|
| | | Value | SE | Value | SE |
| Exponential tumour growth (Eq. (1)), Vehicle | $r$: Intrinsic tumour growth rate (day$^{-1}$) | 0.224 | 0.005 | 0.319 | 0.008 |
| Gompertz tumour growth (Eq. (1)), Vehicle | $r$: Intrinsic tumour growth rate (day$^{-1}$) | 0.0819 | 0.0061 | 0.106 | 0.011 |
| | $K$: Tumour carrying capacity (mm$^3$) | 5460 | 1070 | 7677 | 2307 |
| Logistic tumour growth (Eq. (1)), Vehicle | $r$: Intrinsic tumour growth rate (day$^{-1}$) | 0.305 | 0.004 | 0.423 | 0.019 |
| | $K$: Tumour carrying capacity (mm$^3$) | 1593 | 51 | 2137 | 253 |
| Logistic tumour growth (Eq. (1)), Cisplatin | $r$: Intrinsic tumour growth rate (day$^{-1}$) | 0.355 | 0.011 | 0.327 | 0.019 |
| | $K$: Tumour carrying capacity (mm$^3$) | 1013 | 41 | 939 | 84 |

| | | | | | |
|---|---|---|---|---|---|
| Logistic tumour growth (Eq. (1)), ICB | $r$: Intrinsic tumour growth rate (day$^{-1}$) | 0.376 | 0.024 | 0.297 | 0.014 |
| | $K$: Tumour carrying capacity (mm$^3$) | 1317 | 129 | 5141 | 2386 |
| Logistic tumour growth (Eq. (1)), Cisplatin + ICB | $r$: Intrinsic tumour growth rate (day$^{-1}$) | 0.317 | 0.005 | 0.299 | 0.023 |
| | $K$: Tumour carrying capacity (mm$^3$) | 1055 | 26 | 581 | 57 |
| Logistic tumour growth and cisplatin PK/PD model (Eqs. (2)-(3)), Cisplatin | $k_2$: Linear killing rate of cisplatin (L·mg$^{-1}$·day$^{-1}$) | 8.07 | 2.63 | 63.7 | 4.6 |
| Immune-extended tumour growth model (Eq. (4)), Vehicle | $r$: Intrinsic tumour growth rate (day$^{-1}$) | 1.57 | 0.11 | 2.03 | 0.02 |
| | $p$: Rate of NK cell inactivation by tumour cells (mm$^{-3}$ day$^{-1}$) | 7.49x $10^{-4}$ | 1.63 x $10^{-4}$ | 2.62 x $10^{-4}$ | 7.34 x $10^{-6}$ |
| | $q$: Rate of CD8+T cells inactivation by tumour interaction (mm$^{-3}$ day$^{-1}$) | 2.00 x $10^{-3}$ | 4.91 x $10^{-4}$ | 7.74 x $10^{-4}$ | 2.20 x $10^{-5}$ |
| Immune-extended tumour growth and cisplatin PK/PD model (Eq. (5)), Cisplatin | $r_{C,N}$: Recruitment of NK cells into TME due to cisplatin (L·mg$^{-1}$·day$^{-1}$) | 7.95 | 25.73 | 64.4 | 0.98 |
| | $r_{C,E}$: Recruitment of CD8+ T cells into TME due to cisplatin (L·mg$^{-1}$·day$^{-1}$) | 0 | 419.0 | 97.9 | 9.10 |
| Immune-extended tumour growth model (Eq. (4)), ICB | $q$: Rate of CD8+T cells inactivation by tumour interaction (mm$^{-3}$ day$^{-1}$) | 2.14 x $10^{-3}$ | 1.07 x $10^{-3}$ | 1.41 x $10^{-4}$ | 1.14 x $10^{-4}$ |

**Table S5. Fitted Parameters.** Parameter descriptions, values, standard errors (SE), and which model they are present in for both MP and MPB1 tumours.

*Growth model selection*

We first fit all three tumour growth models of Eq. (1) in the main text to the data from vehicle-treated MP and MPB1 tumours[1]. As expected, each yielded distinct estimates for the growth rate, $r$, and carrying capacity, $K$. Overall, we found all three growth models to be more-or-less consistent with the observed tumour dynamics; however, the logistic and Gompertz models produced nearly indistinguishable fits and more closely captured the experimental data (**Figure S1**). Comparing the AICc and BIC values of each growth model (**Table S1**), $g(T)$, we found the logistic model to best describe the tumour growth dynamics in the MP mice whereas the Gompertz model was preferred for the MPB1 mice, albeit with very small differences in $\Delta$AICc and $\Delta$BIC values between logistic and Gompertzian growth in both cases.

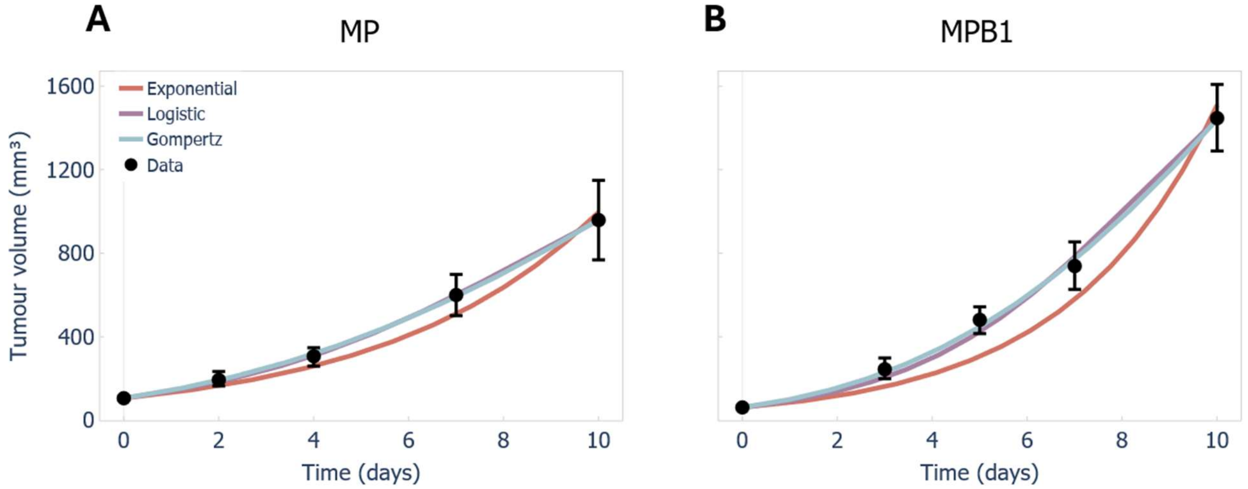


**Figure S1. Predicted MP and MPB1 tumour volume.** Longitudinal tumour volumes for A) MP and B) MPB1 tumours. Fitting results and data favour saturating (i.e., logistic or Gompertz) growth over exponential.

Both logistic and Gompertzian growth laws are special cases of the generalized logistic framework. The choice of an appropriate growth law depends on many factors, such as tumour type. In their study of treatment sequencing in ovarian cancer, Kohandel et al.[2] showed that Gompertzian and generalized logistic growth models lead to essentially the same qualitative conclusions. Similarly, in our model, where the AICc and BIC values for logistic and Gompertzian growth model are comparable, we follow de Pillis et al.[3] and adopt the logistic growth model for the remainder of this work, primarily because of its simpler mathematical structure.

*Tumour growth and immune cells model: Fitting from tumour initiation*

Estimating the parameters in the immune model of Eq. (4) required initial conditions for the numbers of NK cells, $N$, and CD8+ T cells, $E$. Because these quantities were not measured at treatment initiation in Paffenholz et al.[1], we estimated them by simulating tumour growth from tumour initiation, where biologically reasonable assumptions can be made about the immune cell populations in a tumour-free environment.

Specifically, we assumed a tumour volume of $T = 2 \times 10^{-4}$ mm$^3$, $N = N_0/d_N$ cells (homeostatic equilibrium), and $E = 0$ cells at tumour initiation. The time elapsed between tumour initiation and treatment initiation was estimated by solving the logistic growth equation using the parameters fitted to vehicle data. Starting from an initial tumour volume of $T = 2 \times 10^{-4}$

$mm^3$, the estimated times between tumour initiation and the first experimental tumour volume measurements were 43.4 days for MP tumours and 29.9 days for MPB1 tumours.

The experimental time points were then shifted by these estimated intervals so that tumour initiation corresponded to $t = 0$. The model of Eq. (4) was fitted simultaneously to the tumour volume and immune cell data over this period (**Figure S2**), allowing us to estimate the numbers of NK cells and CD8+ T cells at treatment initiation. These estimated immune cell numbers were subsequently used as the initial conditions for all remaining model fits, for which treatment initiation defines $t = 0$.

Additional assumptions were required to fit the immune data. Paffenholz et al.[1] reported only the proportions of NK cells and T cells among CD45+ cells in each sample (their Fig 4C-D), rather than absolute cell counts. Therefore, to estimate the immune cell numbers required by Eq. (4), we assumed both the proportion of CD45+ cells within the sampled cell population and the proportion of CD8+ T cells among the reported T cell population. These assumptions were informed by previously reported estimates in syngeneic mouse models. Taylor et al.[4] measured the percentage of CD45+ cells among live cells in commonly used syngeneic models and reported values ranging from 20-60%, depending on tumour type and time since tumour implementation. Rodriguez et al.[5] reported that approximately 25% of T cells were CD8+ T cells in syngeneic mouse models of ovarian cancer. To assess the sensitivity of the estimated immune cell counts to these assumptions, we considered CD45+ cell proportions of 20%, 40%, and 60% of the total cell population, together with CD8+ T-cell proportions of 20%, 30%, and 40% of the T-cell population.

We selected the assumption that CD45+ cells comprised 60% of the sampled cells and CD8+ T cells represented 30% of the T-cell population, as this combination provided good agreement with the experimental tumour growth data (shown in **Figure S2**). Under these assumptions, the estimated immune cell counts at treatment initiation were $N = 3.82 \times 10^5$ cells and $E = 1.29 \times 10^5$ cells for MP tumours, and $N = 4.03 \times 10^5$ cells and $E = 7.19 \times 10^4$ cells for MPB1 tumours.

Importantly, the estimated initial immune cell counts were relatively insensitive to the specific assumptions. For MP tumours, the different combinations predicted NK cell counts between $3.76 \times 10^5$ and $3.88 \times 10^5$ cells, and CD8+ T cell counts between $1.07 \times 10^4$ and $1.50 \times 10^4$ cells. Similarly, for MPB1 tumours, the combinations yielded NK cell counts between $4.01 \times 10^5$ and $4.07 \times 10^5$ cells and CD8+ T-cell counts between $6.17 \times 10^4$ and $7.77 \times 10^4$ cells. These relatively narrow ranges suggest that the inferred immune cell counts at treatment initiation are robust to choices of the assumed CD45+ and CD8+ T cell proportions.

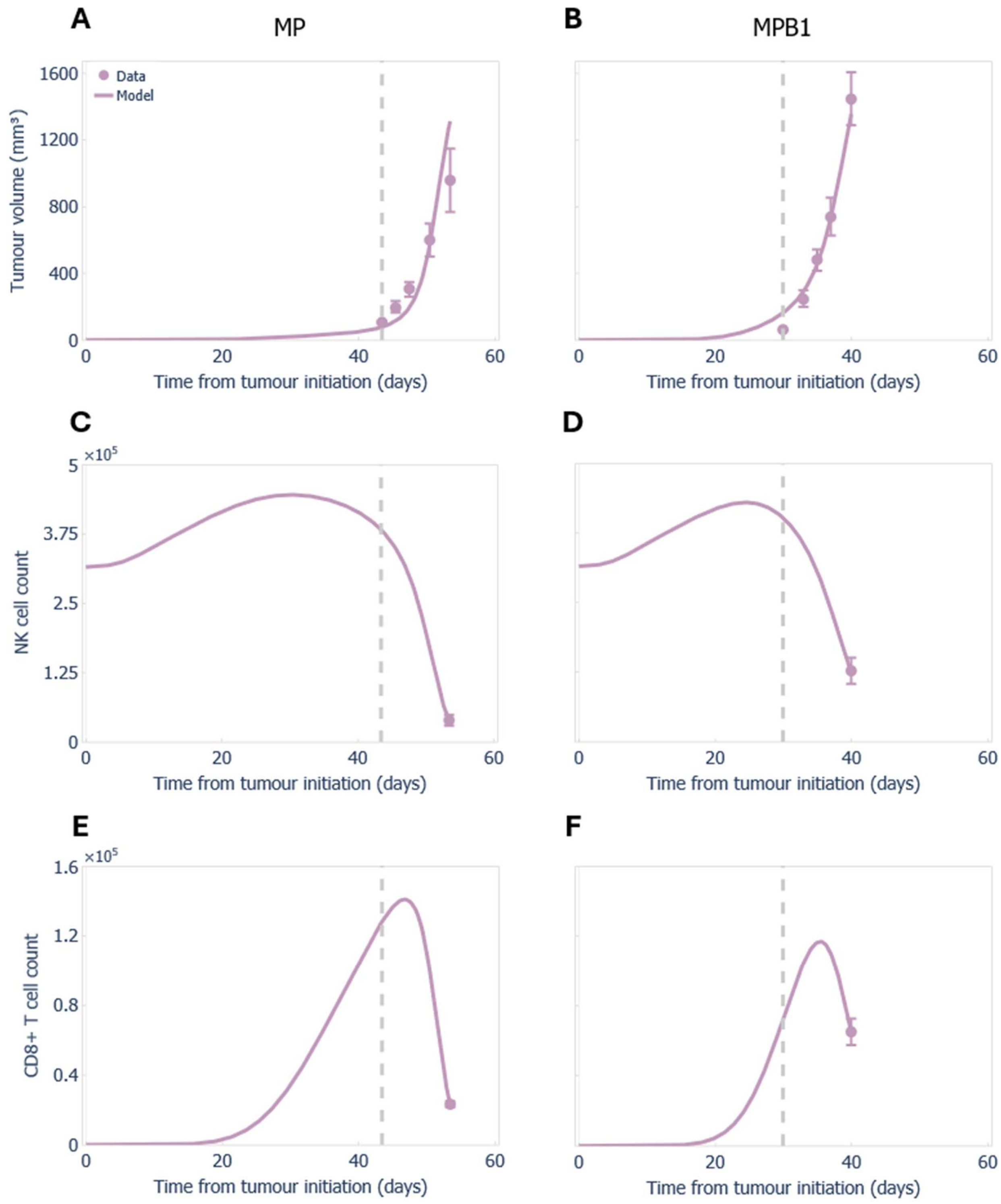


**Figure S2. Tumour-immune model estimates immune cell numbers at treatment initiation.** Predicted tumour growth and immune cell levels from tumour initiation for the tumour-immune extended model (Eq. (4)) fitted to vehicle (untreated) tumour volume measurements (**A-B**), NK cell counts (**C-D**), and CD8+ T cell counts (**E-F**) in MP (**A, C, E**) and MPB1 (**B, D, F**) tumours. Solid lines: model predictions; circles: mean data; error bars: standard error of the mean (SEM), vertical dashed line: treatment initiation.

*Profile Likelihood Results*

The practical identifiability of the estimated parameters was assessed using profile likelihood analysis, as described in the **Methods**. The multiplicative lower and upper bound coefficients used to define the parameter ranges are reported in **Table S9**, and the resulting univariate profile likelihoods are shown in **Figures S3-S6**.

| Parameter | Description and units | $C_L$ | $C_U$ | Reference |
|---|---|---|---|---|
| $r$ | Intrinsic tumour growth rate ($day^{-1}$) | 0.350 | 0.350 | Figures S3 and S4 |
| $K$ | Tumour carrying capacity ($mm^3$) | 0.350 | 1.750 | Figures S3 and S4 |
| $r$ (immune extended model) | Intrinsic tumour growth rate ($day^{-1}$) | 0.200 | 0.200 | Figure S5 |
| $p$ | Rate of NK cell inactivation by tumour cells ($mm^{-3}$ $day^{-1}$) | 0.400 | 1.000 | Figure S5 |
| $q$ | Rate of CD8+T cells inactivation by tumour interaction ($mm^{-3}$ $day^{-1}$) | 0.400 | 1.000 | Figure S5 |
| $r_{C,N}$ | Recruitment of NK cells into TME due to cisplatin ($L \cdot mg^{-1} \cdot day^{-1}$) | 0.175 | 0.175 | Figure S6 |
| $r_{C,E}$ | Recruitment of CD8+ T cells into TME due to cisplatin ($L \cdot mg^{-1} \cdot day^{-1}$) | 0.250 | 0.175 | Figure S6 |

**Table S9. Multiplicative lower and upper bound coefficients defining the range of points for the likelihood profiles.** Lower and upper scaling factors, $\boldsymbol{C_L}$ and $\boldsymbol{C_U}$, respectively, used to define parameter profile likelihood curves (**Figures S3-S6**).

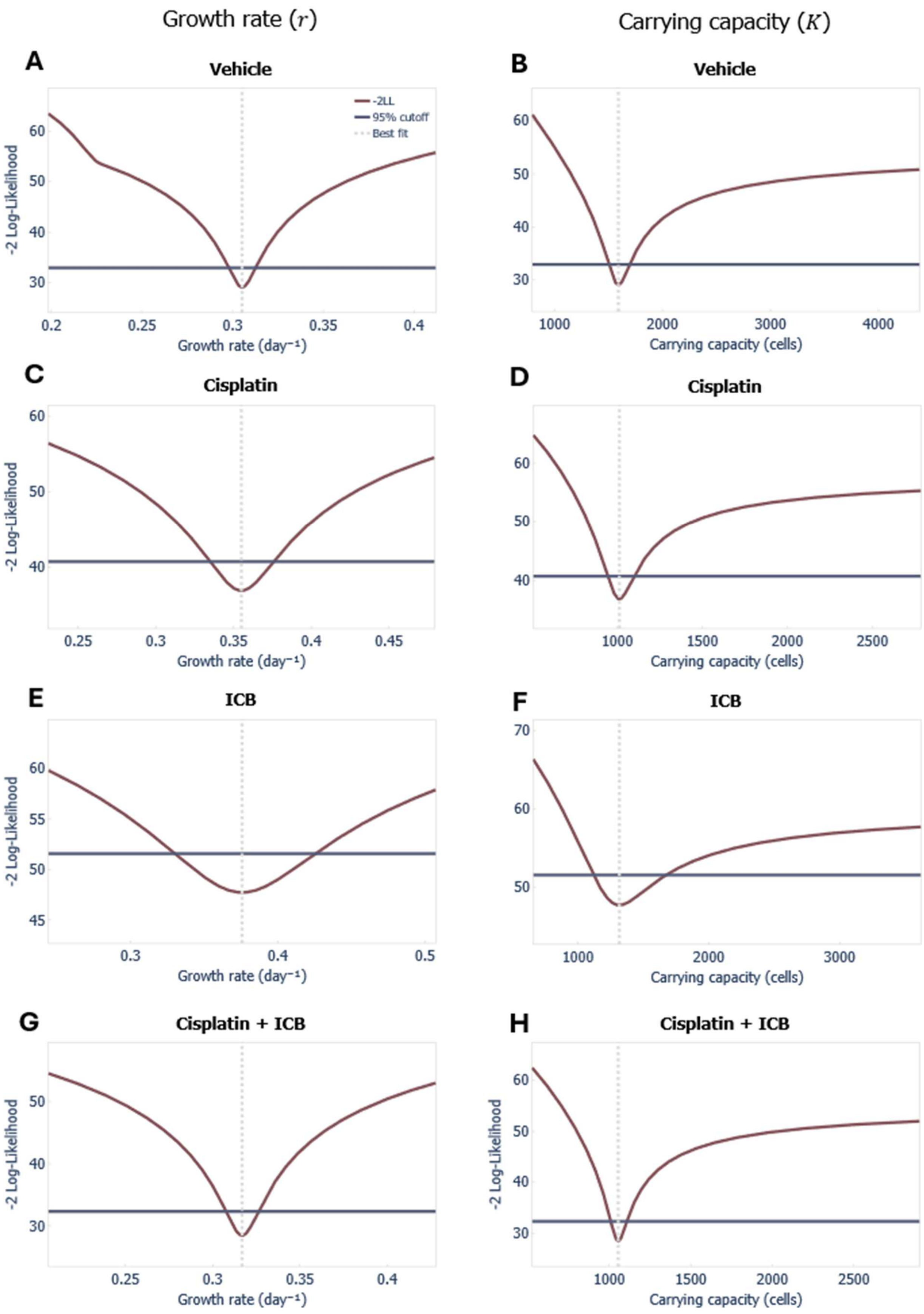

Growth rate ($r$)
Carrying capacity ($K$)
A
Vehicle
-2LL
95% cutoff
Best fit
-2 Log-Likelihood
Growth rate (day⁻¹)
B
Vehicle
Carrying capacity (cells)
C
Cisplatin
D
Cisplatin
E
ICB
F
ICB
G
Cisplatin + ICB
H
Cisplatin + ICB

**Figure S3. Univariate likelihood profiles for parameters estimated from logistic growth models fitted to vehicle, cisplatin, ICB, and cisplatin + ICB treated MP tumours.** Univariate likelihood profiles obtained from fitting the logistic growth model of Eq. (1). Profiles for the growth rate are shown in **A**, **E**, **C**, **G**, and for the carrying capacity in **B**, **D**, **F**, **H**, corresponding to vehicle, cisplatin, ICB, and cisplatin + ICB treatments, respectively. The vertical dashed line indicates the maximum likelihood estimate, and the horizontal solid line indicates the 95% confidence threshold.

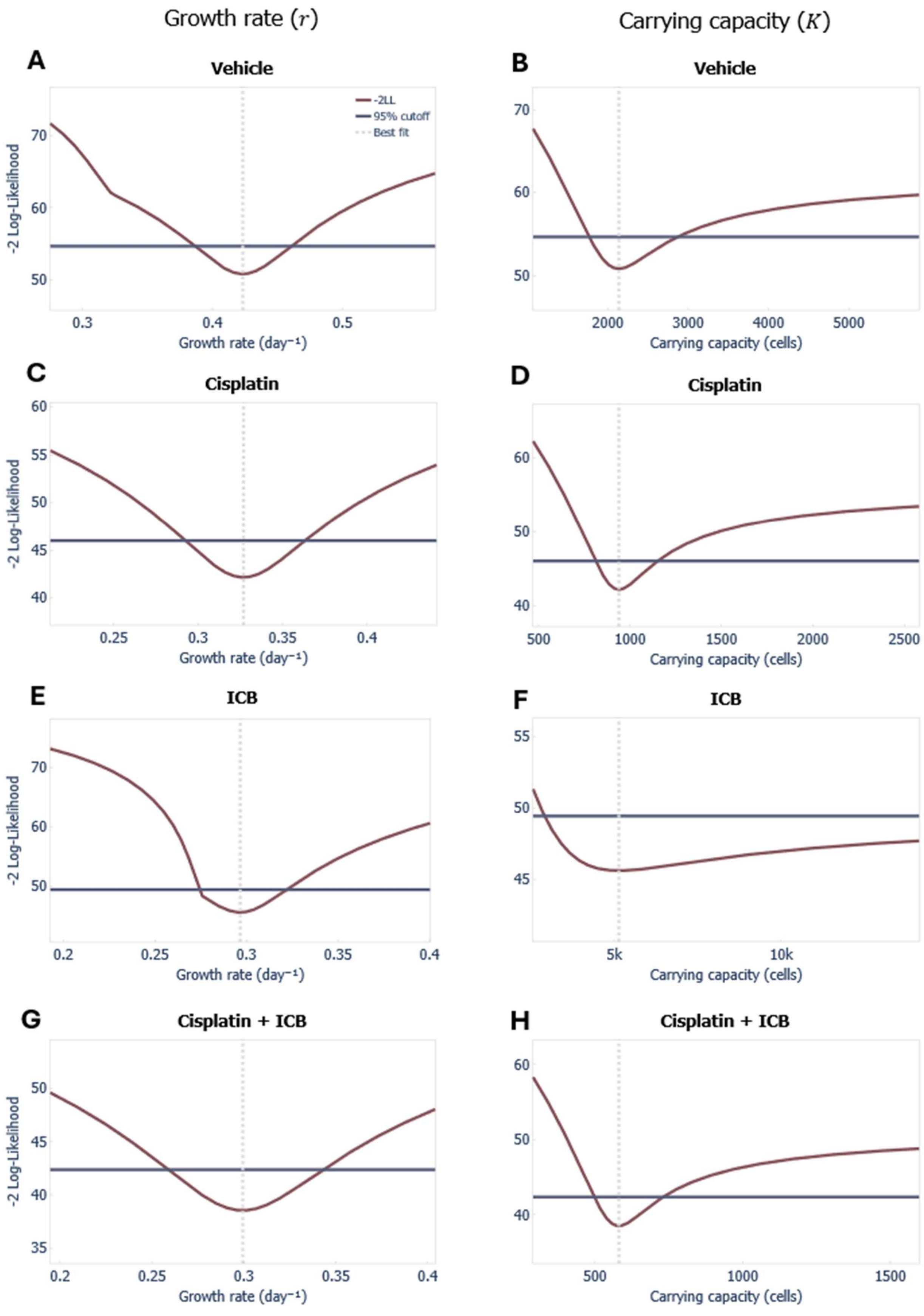

Growth rate ($r$)
Carrying capacity ($K$)
A
Vehicle
-2LL
95% cutoff
Best fit
-2 Log-Likelihood
Growth rate (day⁻¹)
B
Vehicle
Carrying capacity (cells)
C
Cisplatin
D
Cisplatin
E
ICB
F
ICB
G
Cisplatin + ICB
H
Cisplatin + ICB

**Figure S4. Univariate likelihood profiles for parameters estimated from logistic growth models fitted to vehicle, cisplatin, ICB, and cisplatin + ICB treated MPB1 tumours.** Univariate likelihood profiles obtained from fitting the logistic growth model of Eq. (1). Profiles for the growth rate are shown in **A**, **E**, **C**, **G**, and for the carrying capacity in **B**, **D**, **F**, **H**, corresponding to vehicle, cisplatin, ICB, and cisplatin + ICB treatments, respectively. The vertical dashed line indicates the maximum likelihood estimate, and the horizontal solid line indicates the 95% confidence threshold.

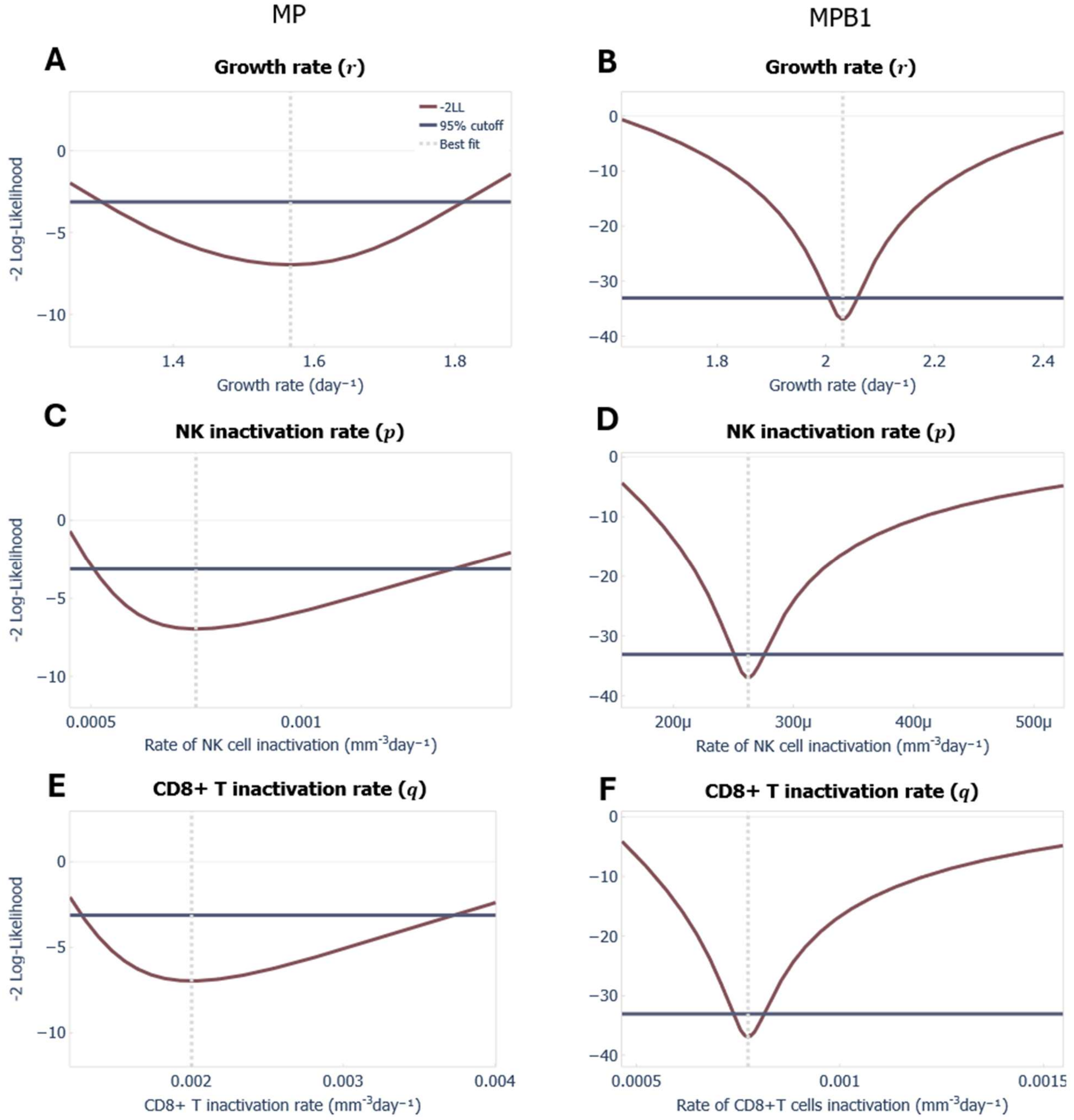


**Figure S5. Univariate likelihood profiles for parameters estimated from the immune extended model fitted to vehicle (untreated) data.** Univariate likelihood profiles obtained from fitting the tumour-immune extended model of Eq. (4). Profiles for the growth rate are shown in **A-B**, NK cell inactivation rate in **C-D**, and CD8+ T cell inactivation rate in **E-F**, with results for MP tumours shown in **A**, **C**, **E** and MPB1 tumours in **B**, **D**, **F**, respectively. The vertical dashed line indicates the maximum likelihood estimate, and the horizontal solid line indicates the 95% confidence threshold.

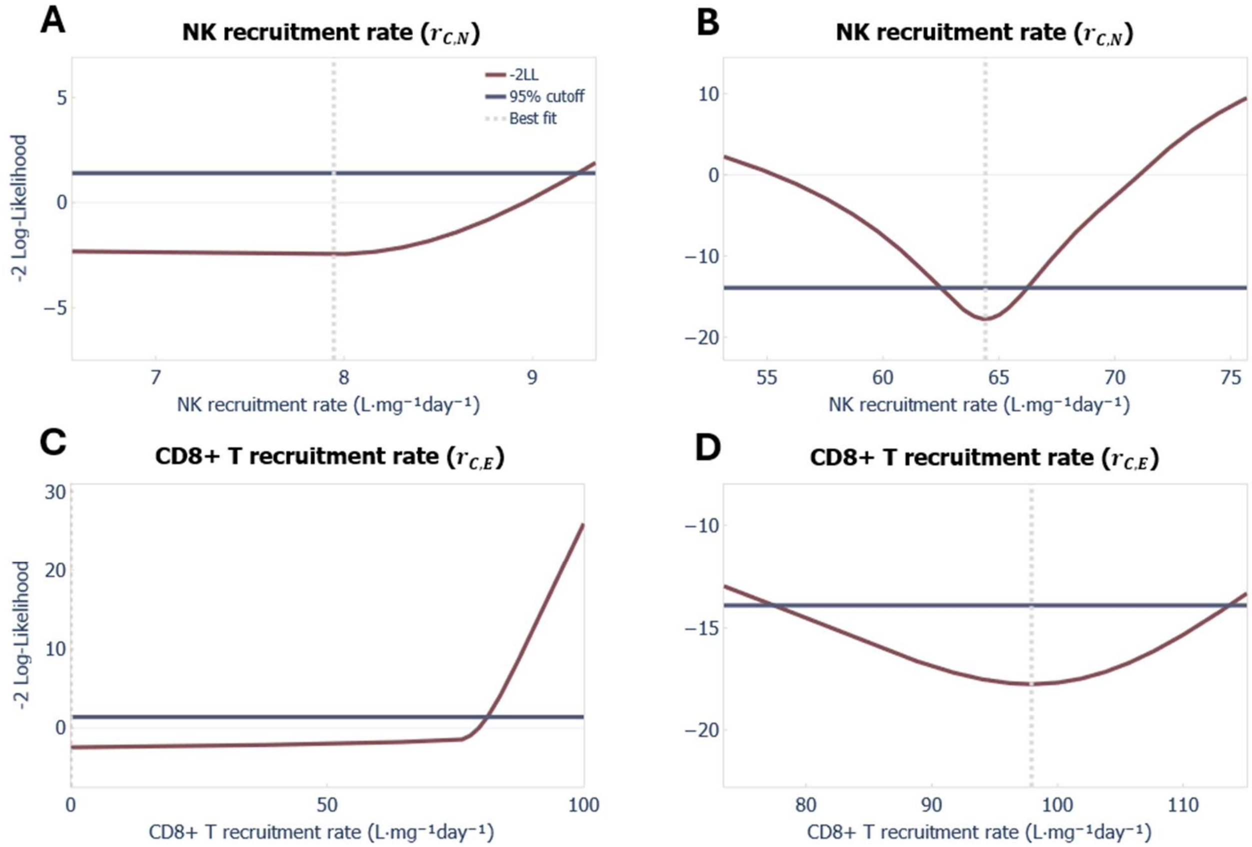


**Figure S6. Univariate likelihood profiles for parameters estimated from the immune extended and cisplatin PK/PD model fitted to cisplatin treated data.** Univariate likelihood profiles obtained from fitting the tumour-immune extended and cisplatin PK/PD model of Eq. (5). Profiles for the NK recruitment rate are shown in **A-B** and CD8+ T cell recruitment rate in **C-D**, with results for the MP tumours shown in **A** and **C** and for MPB1 tumours in **B** and **D**, respectively. The vertical dashed line indicates the maximum likelihood estimate, and the horizontal solid line indicates the 95% confidence threshold.
